# PRDM16 Coordinates Genetic and Epigenetic Programs Governing Chondrogenesis and Chondrocyte Phenotype Specification in the Knee Joint

**DOI:** 10.64898/2026.01.28.701140

**Authors:** Eloise Fadial, Victoria Hansen, Gulzada Kulzhanova, Eliya Tazreena Tashbib, Deeksha Chinta, Alexis Klee, Helen Shammas, Gourango Pradhan, Chia-Lung Wu

**Affiliations:** Dept. of Orthopaedic Surgery and Rehabilitation, Center for Musculoskeletal Research, University of Rochester Medical Center, Rochester, NY 14611; Dept. of Biomedical Engineering, University of Rochester, Rochester, NY 14627; Dept. of Chemical Engineering, University of Rochester, Rochester, NY 14627

**Keywords:** cartilage, bone, Prdm16, Mef2c, epigenetics, CUT&RUN sequencing

## Abstract

Cartilage development and homeostasis require precise regulation by transcriptional and epigenetic networks. PRDM16 is a transcription factor containing zinc finger domains that enable protein-DNA and protein-protein interactions, as well as domains with the capacity for histone methyltransferase activity. However, the detailed molecular mechanisms by which PRDM16 regulates chondrogenesis and chondrocyte identities remain largely unknown. Using our osteochondral lineage-specific, conditional knockout mouse model (*Col2a1Cre;Prdm16^flox/flox^*, Prdm16 cKO), we found that loss of Prdm16 in osteochondral lineage cells delays, but does not fully inhibit, endochondral ossification and bone formation in the knee joint. Furthermore, Prdm16 cKO male mice exhibit comparable OA severity between injured and non-injured joints, suggesting that PRDM16 may exert a chondroprotective function. In our hiPSC-derived chondrocyte model, we observed significantly reduced pellet size and DNA content in cells with modulated PRDM16 expression compared to Control, implying a link between PRDM16 and chondrocyte viability. Integrated analysis of single cell RNA-sequencing and CUT&RUN-sequencing revealed that PRDM16 regulates chondrocyte cell fate decisions by altering chromatin accessibility and DNA binding at promoter/enhancer regions of genes essential for chondrogenesis and chondrocyte hypertrophy. Indeed, PRDM16 governs the expression of key chondrogenic regulators including *SOX9*, *ARID5A*, *SMOC2*, *HAND2*, and hypertrophic driver *MEF2C*. Overall, our results provide evidence that PRDM16 serves as an essential genetic and epigenetic regulator of chondrogenesis and chondrocyte phenotype specification in the knee joint through DNA binding and by modulating H3K4me3 histone mark deposition.

## Introduction

Osteoarthritis (OA) is the leading degenerative joint disease, affecting over 500 million people worldwide^1^. OA is associated with numerous risk factors including aging, sex, obesity, trauma, joint instability, and abnormal loading^2^. The associated abnormal joint loading drives inflammation and promotes cartilage degradation. Due to its progressive nature, OA eventually develops to the point of major loss of joint function^2^. Despite its enormous clinical burden, no disease-modifying osteoarthritis drugs (DMOADs) are currently available, largely due to an incomplete understanding of the genetic and epigenetic mechanisms governing cartilage development and OA pathogenesis.

In our previous work using Bulk RNA sequencing (RNA-seq) to delineate the molecular landscape of chondrogenesis from human induced pluripotent stem cells (hiPSCs), we observed that PR domain containing 16 (*PRDM16*) is upregulated with other essential chondrogenic markers including *ACAN* and *SOX6*, suggesting that PRDM16 may be critical in regulating chondrogenesis^3^. PRDM16 possesses PR domains with the capacity for histone methylation, and zinc finger domains to enable protein-DNA and protein-protein interactions, respectively^4^. PRDM16 has been previously described to have diverse biological functions, including a master regulatory role in promoting brown fat adipogenesis^5^, and involvement in neurogenesis^6^ and cardiovascular development^7^. In humans, microdeletion of the 1p36 chromosome locus encompassing PRDM16 is linked to delayed development, short stature, and orofacial clefting^8^; clinical features characteristic of Pierre Robin anomalad^9^.

Loss of either *prdm3* or *prdm16* has been linked to numerous craniofacial defects including mild hypoplasia of Meckel’s cartilage in zebra fish^10^. During mouse embryonic development, *Prdm16* is expressed in the limb bud mesenchyme at embryonic timepoint (E) 11.5 and remains detectable in the E15.5 mouse knee joint^11, 12^. As the limb develops, *Prdm16* expression is gradually restricted in the mesenchyme and diverted for cartilage condensation^12^. Recent studies have also shown that global knockout (gKO) of *Prdm16* in mice is neonatal lethal and results in abnormal osteogenic and chondrogenic differentiation, while heterozygous KO of *Prdm16* inhibits endochondral ossification in the mouse femoral head^10, 12, 13^. Importantly, PRDM16 is also a top downregulated gene in the subchondral bone of patients with OA^14^. However, the exact molecular mechanisms by which PRDM16 regulates embryonic knee development and adult cartilage homeostasis remain poorly understood.

Here, we hypothesize that PRDM16 is a positive regulator in chondrocyte specification and articular cartilage development. We therefore used our osteochondral lineage-specific, *Prdm16* conditional knockout (*Col2a1Cre;Prdm16^flox/flox^*; Prdm16 cKO) mice, inducible hiPSC-derived chondrogenic models, single-cell RNA sequencing (scRNA-seq), and Cleavage Under Targets & Release Using Nuclease sequencing (CUT&RUN-seq) to define the genetic and epigenetic programs governed by PRDM16 throughout knee joint development.

## Materials and Methods

### Mouse Models and Derivation of Prdm16 cKO mice

This study was carried out in strict accordance with the recommendations in the Guide for the Care and Use of Laboratory Animals of the National Institutes of Health. All animal procedures were approved by the University Committee on Animal Research (UCAR) at the University of Rochester (UR). All mice were maintained under pathogen-free conditions and housed with no more than five animals per cage under standardized light-dark cycle conditions, with *ad libitum* access to food and water. The vivarium was maintained under controlled temperature and humidity. *C57/BL6J* mice were obtained from the Jackson Laboratory (Bar Harbor, ME, USA, #000664). *Prdm16^f^*^lox^ mice (B6.129-*Prdm16^tm^*^1^*^.1Brdp^/J,* #024992, Jackson lab) were crossed with mice whose expression of Cre recombinase was controlled by the *Col2a1* promoter (B6;*SJL-Tg(Col2a1Cre)1Bhr/J)* to generate *Col2a1Cre;Prdm16^flox/flox^* transgenic mice (Prdm16 cKO). Cre-negative littermates were used as controls (WT). Prdm16 cKO was confirmed by Western blot of costal cartilage and bone at E15.5 and E18.5 (**Fig. 1A**).

**Fig. 1:**
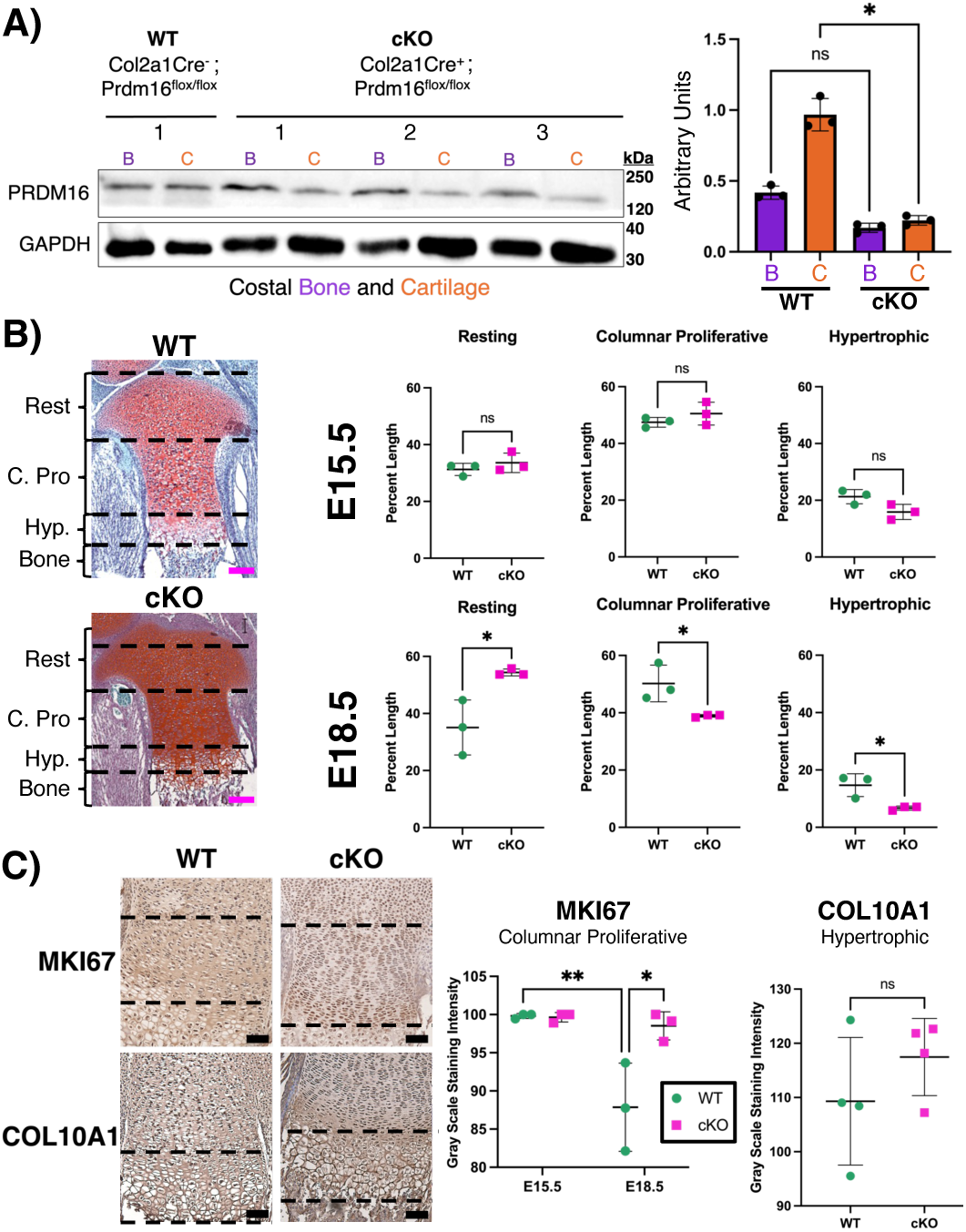
E18.5 cKO mice exhibit symptoms of delayed development compared to WT. (**A**) Western blot of E18.5 costal cartilage and bone (left) and quantified densitometry (right). Each number represents an individual mouse. (**B**) Saf-O/Fast Green staining of E18.5 WT and cKO limbs depicting designated zones for limb measurement analysis. Zones include resting (Rest), columnar proliferative (C. Pro), and hypertrophic (Hyp.) chondrocytes. Pink scale bar is 100µm. Percentage of each zonal length (i.e., individual zonal length / combined total zonal length) in both E15.5 and E18.5 limbs (*p < 0.05). (**C**) IHC of proliferative marker MKI67 (E15.5, E18.5) and hypertrophic marker COL10A1 (E18.5) and respective quantification (*p < 0.05, **p < 0.01). Black scale bar is 50µm. Black dashed lines indicate ROI. Results were analyzed with Student’s Unpaired t-test with Welch correction, or one-way ANOVA as appropriate.

### Destabilization of the Medial Meniscus to Induce Knee OA

At 16 weeks of age, female and male WT and cKO mice underwent destabilization of the medial meniscus (DMM) to induce knee OA in the left hind limbs. Right limbs were used as non-injury control following previously published protocols^15,16^. Hind limbs were harvested 12 weeks post-DMM surgery. The OA severity was quantified through Saf-O/Fast Green staining analyzed by Modified Mankin grading scheme^17^, and synovitis was evaluated with the Krenn synovitis grading scheme^18^.

### Bone Microstructural Analysis

Hind limbs were submitted for µCT scan prior to decalcification to measure bone structural and morphological changes (Scanco Medical VivaCT 40 cone-beam CT, with 10.5 µm isotropic cubical voxels). 1000 projections were generated over 180°, 300msec integration time, and reconstructed with proprietary Scanco algorithms. Scanco evaluation software V6.5 was used to analyze µCT bone parameters. Trabecular bone and cortical bone were measured in the tibia and femur of all samples. The bone volume over total volume ratio (BV/TV) and bone mineralization density (BMD) in the region of interest (ROI) were quantified. These measurements were plotted in GraphPad. Results were analyzed with Student’s Unpaired t-test with Welch correction or One-Way ANOVA with Tukey’s post-hoc as indicated.

### Histological Analysis

#### Embryonic

Mouse hind limbs were harvested at E11.5, E13.5, E15.5, and E18.5. Hind limbs were fixed with in 10% Neutral Buffered Formalin (NBF) for either 3 hours (E11.5, E13.5) or 24 hours (E15.5, E18.5) at room temperature (RT). Limbs were then rinsed with PBS and stored in 70% ethanol at 4°C until processed for paraffin embedding by the UR CMSR Histology Core. Embedded tissues were sectioned via microtome at 7µm thickness and stained by Safranin O (Saf-O)/Fast Green following standard protocols^,19, 20^. Tissues were also stained with Harris Hematoxylin. Chondrocyte zonal lengths were measured in ImageJ as a percentage of total length and plotted in GraphPad Prism (Version 10.5.0). Results were analyzed with Student’s Unpaired t-test with Welch correction.

#### Post-Natal

For post-natal mice, hind limbs were harvested at 4-weeks, 12-weeks, 52-weeks, and 12-week post-DMM surgery (i.e., 28-weeks of age). Hind limbs were fixed in 10% NBF at RT for 48 hours and then rinsed with PBS and stored in 70% ethanol at 4°C until ready for decalcification. CalEx (Fisher CS510-1D) decal solution was applied for 72 hours at 4°C to decalcify tissues. Tissues were rinsed for 5 minutes with 1x PBS (3 times), diH2O (3 times), 50% EtOH (1 time), and 70% EtOH (1 time). Tissues were stored in 70% EtOH at 4°C or directly processed for paraffin embedding by the UR CMSR Histology Core. Embedded tissues were sectioned via microtome at 8µm thickness and stained by Saf-O/Fast green and Harris Hematoxylin, or Hematoxylin and Eosin/Orange G stain following standard protocols^20, 21^.

### Immunohistochemical Analysis

Primary antibody (Ab) staining was performed with either PRDM16 polyclonal Ab (ThermoFisher Scientific, PA5-20872; 1:500 dilution), Ki67/MKI67 Ab (Novus Biologicals, NB500-170; 1:250 dilution), COL10A1 Ab (Abcam, ab49945; 1:500 dilution), or COL2A1 Ab (ThermoFisher Scientific, PA126206; 1:200 dilution) for mouse tissues and Collagen X Ab (ThermoFisher Scientific, PA5-116871; 1:200 dilution) for human tissues. ImmPRESS polymer detection kit (Vector Laboratories, MP-7404-50) was used as the secondary antibody. ImmPACT DAB substrate kit solution (Vector Laboratories, SK-4105) was applied, and color change was monitored. Sections were counterstained with Hematoxylin QS (Vector Laboratories, H-3404). Tissue slides were then dehydrated and cover slipped with Optic Mount I (Mercedes Scientific, MER 7724). Counts of percent-stained nuclei or percent-stained area were performed, as appropriate using ImageJ and analyzed in GraphPad.

### Generation of PRDM16 Overexpressed, Knockdown, and Knockout hiPSC lines

Cells with doxycycline inducible knockdown (KD) or overexpression (OE) of PRDM16 were generated from STAN61i-164-1 hiPSC line (WiCell Research Institute): Non-targeted control hiPSC line (SMARTvector Inducible Lentiviral Controls, promotor: hEF1a, Horizon, #VSC6573), PRDM16 OE hiPSC line (pLVX-TetOne-hPRDM16-Puro vector, Clontech, #631894), and PRDM16 KD hiPSC line (SMARTvector Inducible Lentiviral shRNA with a set of 3 shRNAs: Clone Id: V3IHSHER_5736860, Clone Id: V3IHSHER_6066299, and Clone Id: V3IHSHER_5195528, promotor: hEF1a, reporter: TurboRFP, Horizon)^22^. We also used CRISPR/Cas9-mediated genome editing to knockout (KO) PRDM16 in STAN61i-164-1 hiPSCs. Transfected cells were expanded and plated at low density for single cell clonal isolation. After 10–14 weeks, the resulting 8 clones were manually selected and expanded for Western blot screening (**Supp. Fig. 1A**). Clone 4 (red box) was used for chondrogenesis and downstream CUT&RUN-seq.

### hiPSC Chondrogenesis

Non-target control (Control), KD, OE, or KO hiPSCs underwent mesodermal differentiation followed by chondrogenic differentiation for 28 days following our previously established protocols^22, 23^. Since PRDM16 expression was previously observed to be increased at the chondroprogenitor (C_P_) stage of differentiation^24^ (**Supp. Fig. 2**), doxycycline (3 µg/ml) was administered at this stage to induce OE or KD expression of PRDM16. Doxycycline treatment was continued throughout the C_P_ stage and chondrogenic differentiation (i.e., pellet culture).

### Human Cell Culture Sample Harvest

Chondrogenic pellets were rinsed once with DPBS (Gibco, #15203147) and then processed for biochemical assay, histology, or scRNA-seq.

### Biochemical Assay and Quantification

Individual pellets were digested in 400µL papain solution^25^ for 24 hours at 50°C. Digested pellets in papain were stored at -20°C until needed for 1,9-dimethylemethylene blue (DMB) or PicoGreen assays, respectively. DMB dye (Sigma, #341008) was prepared by adding 2.0g Sodium Formate (Sigma, 71539-500G) to 5mL 100% EtOH. 21mg of 1,9-dimethylemethylene blue was dissolved into this solution. Total solution was poured into 500mL diH_2_O. Formic acid was used to adjust pH to 3.0. Samples and C-4-S standard (Sigma, C9819-5G) were thawed to RT. Standards were diluted on a range from 0 to 60 µg/mL. 30µL of samples were plated in duplicates with 125µL DMB dye and run through a microplate reader. PicoGreen assay samples and λDNA standard were thawed to RT. Standards were diluted on a range from 0 to 1000 ng/ml. Samples were plated in duplicates using the Quant-iT™ PicoGreen™ dsDNA Assay Kit (ThermoFisher Scientific, P7589) and run through a microplate reader. Data were analyzed in GraphPad.

### hiPSC-Derived Chondrogenic Pellets Histological and Immunohistochemical Staining

Pellets were fixed in 10% NBF for 24 hours at RT. Pellets were then transferred to 70% EtOH and incubated for 24 hours at 4°C prior to paraffin processing. Paraffin-embedded tissues were sectioned via microtome at 7µm thickness and stained by Saf-O/Fast Green, anti-PRDM16 Ab, and anti-COL10A1 Ab as previously described.

### scRNA-seq and CUT&RUN-seq Sample Preparation

Pellets were transferred to digestion solution (0.4% type II collagenase, 5% FBS, 1% P/S in DMEM/F12, warmed to 37°C) and placed in a rotator incubator for 40 min - 1 hour until all pellets were digested. At 30 minutes of digestion, any remaining tissues were crushed to decrease digestion time. Once pellets were digested, neutralization media (20% FBS, 1% P/S in DMEM/F12) was administered in a greater volume than the digestion media. Cells were centrifuged at 300x g for 5 min and supernatant was aspirated. Cells were resuspended in neutralization media, counted with the Cell Countess (ThermoFisher Scientific), and filtered through 40µm Falcon strainer. Cells were centrifuged following same parameters, supernatant was aspirated, and cells were resuspended in freezing media (80% FBS, 10% DMSO, 10% DMEM/F12). Cryovials of cells were placed in Mr. Frosty (ThermoFisher Scientific, 5100-0036) and stored at -80°C for 24-72 hours before being transferred to -150°C for storage until ready for submission.

### scRNA-seq

Samples with greater than 93% viability were submitted to UR Genomics Research Center for scRNA-Seq library preparation via Chromium Next GEM Single Cell 3′ GEM, Library and Gel Bead Kit V3.1 (10x Genomics), per the manufacturer’s recommendations. Post-capture viability ranged from 60-75%. Samples were loaded on a Chromium Single-Cell Instrument (10x Genomics, Pleasanton, CA, USA) to generate single-cell Gel Bead-in-Emulsions (GEMs). GEM reverse transcription (GEM-RT) was performed to produce barcoded, full-length cDNA from poly-adenylated mRNA. After incubation, GEMs were broken, the pooled GEM-RT reaction mixtures were recovered, and cDNA was purified with silane magnetic beads (Dynabeads MyOne Silane Beads, ThermoFisher Scientific). The purified cDNA was further amplified by PCR to generate sufficient material for library construction. Enzymatic fragmentation and size selection were used to optimize the cDNA amplicon size and indexed sequencing libraries were constructed by end repair, A-tailing, adaptor ligation, and PCR. The final libraries contain the P5 and P7 priming sites used in Illumina bridge amplification. Sequence data was generated using the Illumina NovaSeq 6000 platform.

### Bioinformatic analyses

#### Quality Metrics Analysis and initial subgrouping of major cell types

Pre-processed aligned datasets of cells from Cell Ranger were loaded into RStudio to create Seurat objects (Seurat V5.0.1)^26^. Later, a quality metrics analysis was performed to exclude low-quality cells or cells likely undergoing apoptosis. Cells with the highest and lowest RNA features as well as with high percentage of mitochondrial genes (>15%) were excluded from further downstream analyses (**Supp. Table 1)**. Individual datasets were later normalized using SCTransformation^27^. Unsupervised clustering was performed to identify various cell populations and annotated based on expression of conventionally used chondrocyte population markers from prior literature. For example, *HMGB2^+^* cells were annotated as proliferative chondrocytes^28–30^.

#### Integration of Control, KD, and OE PRDM16 pellets

To integrate scRNA-seq datasets across the treatment conditions (i.e., Control, KD, and OE), anchor-based integration was performed following Seurat tutorials. Using the *SelectIntegrationFeatures* command, the top 2000 variable features were identified. Seurat objects were normalized using SCTransform and then applied as the input to find anchors (*FindIntegrationAnchors).* Anchors were used to integrate all datasets using the *IntegrateData* function. *RunPCA* analysis was performed for linear dimensionality reduction and an Elbow Plot was used to identify the percentage of variance explained by each principal component and to determine the number of dimensions. Non-linear dimensional reduction of integrated treatment groups was performed by Uniform Manifold Approximation and Projection (UMAP)^31^. To further determine heterogeneity of cell composition, *FindNeighbors* and *FindClusters* were performed and a resolution of 0.5 was used to identify 11 conserved clusters across all treatment groups (**Fig. 5D**). Next, the *FindConservedMarkers* function was used to determine defining markers of the conserved cell populations. The top 50 upregulated genes of each identified conserved cell population were confirmed via the *VlnPlot* and *FeaturePlot* functions. Next, we identified the condition-specific differentially expressed gene sets for each cell cluster using *FindMarkers*. Differentially expressed genes were determined with p<0.05. The percentage of cells in each cluster with respect to total cells per treatment group was also quantified (**Supp. Table 2**). Note that because (1) PCNA^+^/HMGB2^+^ (Cluster 7) and PLK1^+^/HMGB2^+^ (Cluster 8) proliferative chondrocytes did not differ significantly among Control, OE, and KD treatment conditions, (2) COL1A1^high^/ACTA2^+^ cells (Cluster 4) represent perichondrial cells (as we previously identified^3^), and (3) SOX2^+^ cells correspond to an off-target neuronal population^24^, only Clusters 0, 1, 2, 3, 5, 6, and 10 were further subset and re-clustered for downstream analyses, focusing on the role of PRDM16 in chondrogenesis. This process resulted in 5 distinct chondrocyte subpopulations (**Fig. 5E & Supp. Table 3**). To elucidate the functional differences between chondrocyte clusters, and the functional roles of differentially expressed genes (DEGs) within the same cluster under different treatment conditions (e.g., KD vs. Control), Gene Ontology (GO) analysis and gene set enrichment analysis (GSEA) were performed and visualized using *clusterProfiler* R package^32^.

#### Differentiation trajectory analyses

Several pseudotime analyses including RNA velocity, random walk, Palantir, CellRank 2, and Monocle 3 were applied to elucidate how modulated PRDM16 expression influences chondrocyte cell fates. For RNA velocity, we used the *scVelo* toolkit^33^. In brief, Seurat objects of PRDM16 Control, KD, and OE pellets were converted into AnnData objects in a Python environment (V3.6). Loom files from each condition were generated from FASTQ files using *loompy* (V3.0; RRID:SCR_016666), following instructions with index from Kallisto V0.50 (Homo_sapiens.GRCh38.dna.primary_assembly.fa.gz)^34^. The “dynamical model” was used for RNA velocity estimation. We next applied CellRank 2^35^ to combine RNA velocity with expression similarity in high dimensions using the *VelocityKernel* (vk) and *ConnectivityKernel* (ck) functions. This integration helped smooth noise from the RNA velocity. The default combined_kernel = 0.8*vk + 0.2*ck was used for each condition. Cellular dynamics on the Markov chain defined by the transition matrix (i.e., random walks) were simulated using the *vk.plot_random_walks* function (max_iter=500, seed=0, and s=500). We next applied CellRank 2 and Palantir algorithms (V1.4.1)^36^ to determine chondrocyte cell fate probabilities. Initial differentiation sites were selected based on the RNA velocity and random walk results. For all conditions, SOX5^high^/SOX6^high^ chondroprogenitors were identified as the initial state, while distinct terminal states were predicted by the algorithms. For example, UCMA^+^ chondrocytes and ANGPTL4^high^ chondrocytes were predicted as two terminal states in the Control pellets. Palantir pseudotime was performed using *palantir.core.run_palantir* with num_waypoints = 1000. Expression trends with 95% confidence of *PRDM16*, *SOX6*, *ANGPTL4*, *MT2A*, and *IGFBP5* for each fate lineage, were visualized using Palantir pseudotime as the X-axis. Monocle 3 was also used for differentiation trajectory analyses^31, 37–39^. Size factors were adjusted for best fit per treatment group (Resolutions: Control=0.006, KD=0.004, OE=0.002). The functions *learn_graph* and *order_cells* were used to construct differentiation trajectories and order cells along pseudotime.

#### CUT&RUN-seq

CUT&RUN-seq was performed as previously described^40^. Briefly, single-cell suspension solution from digested d28 chondrogenic pellets (∼ 200,000 cells/sample) with different conditions (i.e., WT or PRDM16 KO) were incubated on activated Concanavalin A conjugated paramagnetic beads following the manufacture’s instruction (CUTANA™, EpiCypher). Cell-bound beads were resuspended in antibody buffer and incubated in the corresponding antibodies: 1) IgG control (EpiCypher, 13-0042), 2) PRDM16 (ThermoFisher Scientific, PA520872), 3) H3K9me1 (EpiCypher, MA5-33385), or 4) H3K4me3 (EpiCypher, 13-0060) were added to samples and incubated overnight with rotation at 4°C. The following day, cells were washed 3 times in Digitonin Buffer wash solution (20 mM HEPES, pH 7.5; 150 mM NaCl; 0.5 mM Spermidine; Complete-Mini Protease Inhibitor tablet (Roche Diagnostics; 0.01% digitonin). Cells were then incubated with pAG-MNase (EpiCypher) for 10 minutes at room temperature. 100 mM CaCl_2_ was then added to each sample and incubated at 4°C for 2 hours. This reaction was stopped with the Stop Buffer (340 mM NaCl, 20 mM EDTA, 4 mM EGTA, 0.05% Digitonin, 0.05 mg/mL glycogen, 5 µg/mL RNase A) for 10 minutes at 37°C. DNA fragments were then purified and released. Next, DNA samples were quantified using Qubit dsDNA assay (ThermoFisher Scientific, Waltham, MA) with up to 5ng of DNA used as the input for NEBNext Ultra II (New England Biolabs, Ipswich, MA) library prep, per manufacturer’s instructions with minor modifications for the CUT&RUN sample type. Briefly, following end repair, Illumina-compatible adapters were ligated to CUT&RUN DNA fragments and a 1.1X Ampure bead purification was performed to retain fragments >150bp. Libraries were amplified with dual-index Illumina-compatible primers with the following parameters: 45s at 98°C, 14 cycles of 15s at 98°C followed by 10s at 60°C, and final extension at 72°C for 1 minute. Amplified libraries were purified with a final 1.1X Ampure bead clean-up, quantified by Qubit dsDNA assay, and qualitatively assessed by Agilent Tapestation. Dual-indexed libraries were pooled on an equimolar basis and sequenced on Illumina’s NovaSeq 6000. HOMER (Hypergeometric Optimization of Motif EnRichment; http://homer.ucsd.edu/homer/download.html) was used for peak calling in the Anaconda environment. The binary alignment map (Bam) files were used to generate tag directories. We then used *findPeaks*, *-style factor* for PRDM16, and *-style histone* for H3K9me1 and H3K4me3 to generate regions.txt files. Annotated peaks were aligned to hg38 reference genome. The regions.txt files were converted to BedGraph files. Coverage heatmaps and profile plots demonstrating the differences 1.5 kb upstream and downstream of centered peaks of PRDM16 DNA binding, H3K4me3, and H3K9me1 across the genome for WT and PRDM16 KO cells were visualized by *deepTools2*^41^ with bin size = 10. The genome track plots for selected genes were generated by *pyGenomeTracks*^42^.

#### Bulk RNA-seq

PRDM16 expression over the course of hiPSC chondrogenesis was examined using our previously published work (GEO GSE160786)^24^. Briefly, un-normalized gene counts were read using DESeq2 R package with *DESeqDataSetFromMatrix*, as instructed by the package tutorial^43^. Genes that were expressed by fewer than 10 cells were excluded. Next, we used *DESeq* and *results* functions, which implement the Wald test in DESeq2, to determine the differentially expressed genes (DEGs) between two consecutive differentiation stages. In this process, the estimation of size factors (i.e., controlling for differences in the sequencing depth of the samples), the estimation of dispersion values for each gene, and fitting a generalized linear model were performed. Gene counts were averaged from three hiPSC lines. To observe the temporal expression of a given gene for each hiPSC line, the count matrix was first regularized-logarithm transformed via *rlog*. We then used *plotCounts* in DESeq2 to visualize the expression pattern of the gene. To evaluate PRDM16 expression levels between healthy and OA cartilage in humans, we applied the datasets from a recent study^44^ (GEO GSE114007) using the *GEO2R* function of the NCBI GEO website. Normalized PRDM16 expression was then visualized in GraphPad Prism.

### Statistical analysis

Detailed statistical analyses are described in the figure captions. Analyses were performed using GraphPad Prism, with significance reported at the 95% confidence level. The default statistical analyses embedded in above-mentioned R packages and Python toolkits were used for bulk RNA-seq, scRNA-seq, and CUT&RUN-seq datasets, accordingly.

## Results

### Loss of *Prdm16* in mice leads to altered chondrocyte zonal length at E18.5

To confirm that PRDM16 was knocked out in a cartilage-specific manner, costal cartilage and bone were harvested from WT and Prdm16 cKO mice at E18.5. Western blot analysis revealed no statistically significant difference in PRDM16 expression in the costal bone between WT and Prdm16 cKO mice. However, costal cartilage of Prdm16 cKO mice had significantly reduced PRDM16 expression compared to WT mice (**Fig. 1A**). This result demonstrates that the Prdm16 cKO model effectively targets PRDM16 expression in the cartilage, but not in other tissues. Although minimal expression of PRDM16 remained detectable in the costal cartilage, this likely resulted from possible contamination of other tissues, such as muscles, due to the challenges in precisely removing unwanted tissues in the small embryonic rib cage at E18.5. No difference in the percent length of the resting, columnar proliferative, or hypertrophic zones was observed between WT and Prdm16 cKO mice at E15.5 (**Fig. 1B**). However, by E18.5, Prdm16 cKO mice exhibited a significantly increased percent length of the resting chondrocyte zone and significantly reduced percentages of columnar and hypertrophic chondrocyte zones as compared to WT (**Fig. 1B**). Immunohistochemical staining (IHC) revealed comparable expression of proliferative marker MKI67 in the columnar proliferative chondrocytes between WT and Prdm16 cKO mice at E15.5 (**Fig. 1C**). However, at E18.5 Prdm16 cKO mice had significantly higher MKI67 expression in the columnar proliferative cells compared to WT. Interestingly, while WT mice exhibited decreased MKI67 in columnar proliferative chondrocytes over time, the expression levels of MKI67 in the columnar proliferative chondrocytes of Prdm16 cKO mice remained comparable between E15.5 and E18.5 (**Fig. 1C, left**). Furthermore, E18.5 cKO mice exhibited a significant loss of staining for MKI67 in the perichondrium (**Supp. Fig. 3**). IHC of COL10A1 in E18.5 limbs revealed that COL10A1 expression levels in the hypertrophic zone were comparable between cKO and WT mice (**Fig. 1C, right**).

### No apparent differences were observed in cartilage Saf-O/Fast Green staining and COL2A1 IHC between WT and Prdm16 cKO 12-week-old mice

To understand the role of Prdm16 in postnatal cartilage development and homeostasis, knee joints from both 4-week (adolescent) and 12-week-old (adult) WT and cKO mice were analyzed histologically in a sex-specific manner, although few differences were observed between female and male mice (**Fig. 2A & Supp. Fig. 4**). Despite cKO of PRDM16, no apparent difference was observed in Saf-O/Fast Green staining between WT and cKO mice at either 4- or 12-weeks (**Fig. 2A**). COL2A1 and COL10A1 expression levels also appeared similar between cKO and WT mice. (**Fig. 2A&B)**. No statistically significant difference in body weight was observed between WT and cKO 4- or 12-week-old mice; however, male 12-wk-old cKO mice have a trend of reduced weight compared to WT (**Fig. 2C**).

**Fig. 2:**
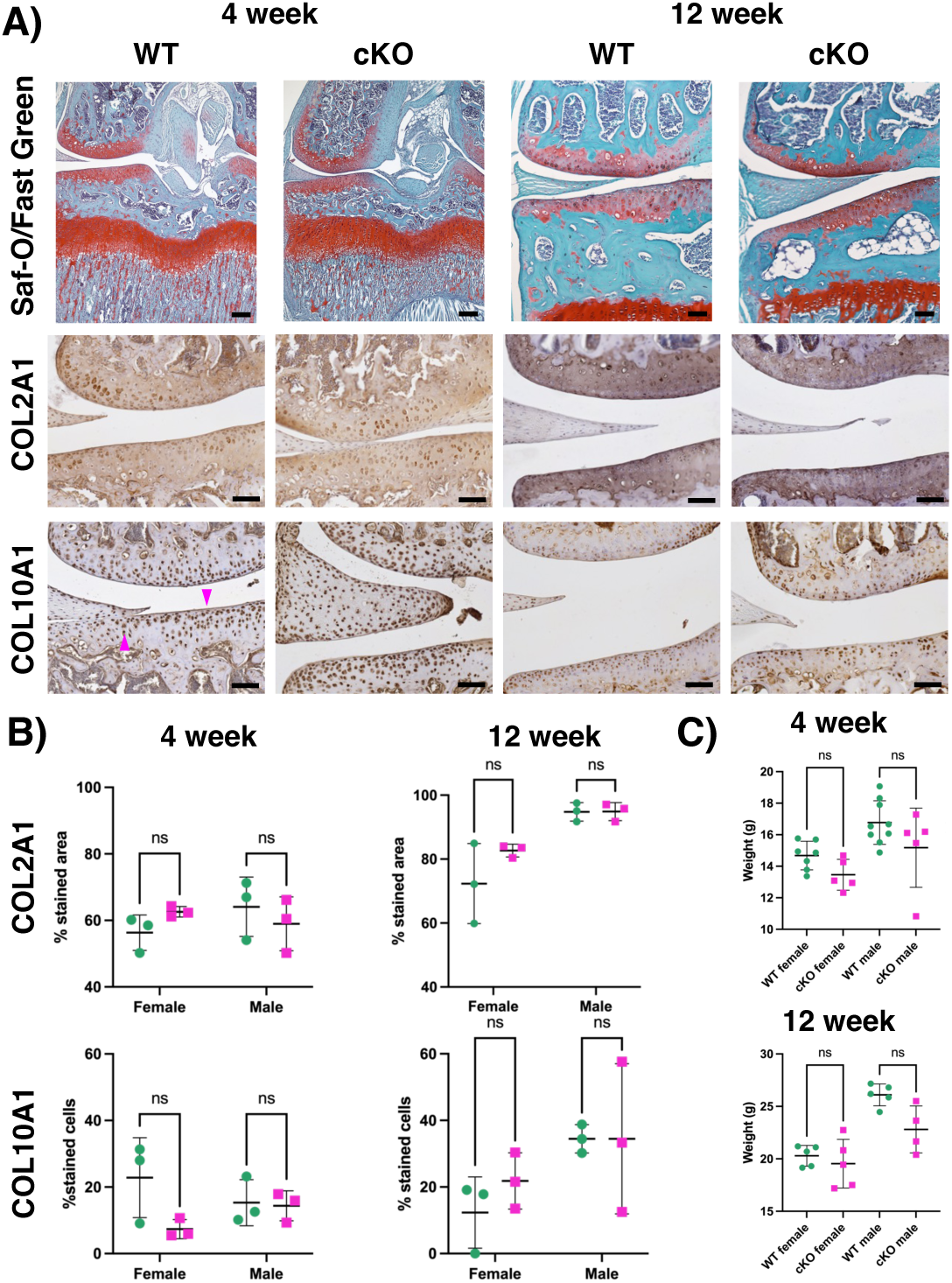
Histological analyses and body weight of 4- or 12-wk-old male PRDM16 WT and cKO mice. (**A**) Histological staining of knee joints: Saf-O/Fast Green, COL2A1, and COL10A1. Black scale bar = 50µm. Pink arrowhead indicates unstained cells. (**B**) No significant difference in COL2A1 or COL10A1 was observed between WT and cKO mice. (**C**) Weights of mice at day of sacrifice. Student’s Unpaired t-test with Welch correction within same sex. ns = nonsignificant.

### µCT revealed decreased knee BV/TV and delayed meniscus ossification in Prdm16 cKO mice

Four distinct regions of ossification were identified and selected for µCT analysis (**Fig. 3A, left**). Both female and male 4-week-old Prdm16 cKO mice exhibited a decreasing trend of knee BV/TV and significantly decreased BV/TV of the ossified medial meniscus compared to WT. Interestingly, 4-wk-old cKO mice of both sexes failed to develop the ossified medial meniscus (**Fig. 3A**; white arrow). The absence of the ossified medial meniscus in the Prdm16 cKO mice was confirmed with BMD analysis (**Supp. Fig. 5**). Additionally, a significant decrease in BV/TV of the ossified patella was observed in 4-week-old male Prdm16 cKO mice compared to WT. (**Fig. 3B**). Interestingly, both female and male 12-wk-old Prdm16 cKO mice maintained significantly decreased BV/TV and BMD of the ossified medial meniscus compared to WT, although other bone parameters were recovered with no significant difference observed between WT and Prdm16 cKO mice (**Fig. 3C & Supp. Fig. 5**).

**Fig. 3:**
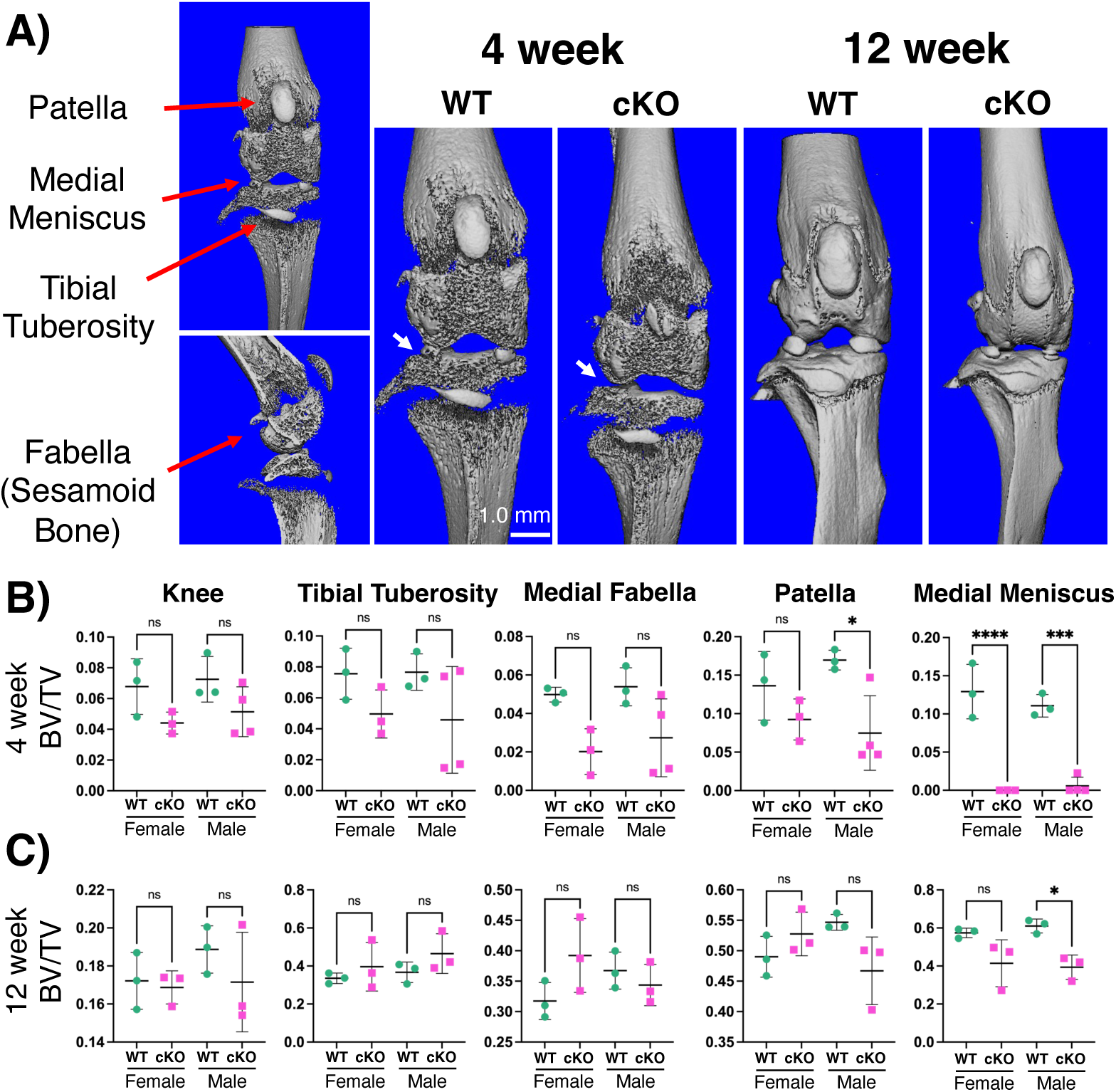
µCT images and analysis of postnatal limbs reveals reduced ossification of medial meniscus in cKO mice vs. WT. (**A**) Regions of interest for analysis (left, red arrows) and images from µCT male 4- and 12-week-old mice (right). (**B**) 4-week-old cKO mice exhibit a trend toward reduced BV/TV across the total knee, tibial tuberosity, and medial fabella relative to WT mice. Male cKO mice further exhibited significantly decreased BV/TV within the patella and medial meniscus compared to WT. White arrow highlights ossified medial meniscus (WT) or lack thereof (cKO). (**C**) 12-week-old mice BV/TV reveals delayed ossification of male cKO medial meniscus. Student’s Unpaired t-test with Welch correction within the same sex (*p < 0.05, ***p < 0.001, ****p < 0.0001).

### Non-surgery joints of male Prdm16 cKO mice exhibit comparable OA severity to DMM-induced OA joints of WT and cKO mice

To further elucidate how PRDM16 expression changes in response to cartilage injury, we surgically induced OA via DMM surgery in both WT and cKO female and male mice (**Fig. 4A**). DMM male mice exhibited symptoms of OA by 12-wks post-surgery including loss of cartilage matrix quality (i.e., loss of Saf-O staining) in the articular cartilage, and osteophyte formation (**Fig. 4B**). Additionally, some male cKO mice exhibited loss or clefts in the articular cartilage following DMM surgery (**Fig. 4B**, bottom right). Interestingly, non-surgery joints of Prdm16 cKO male mice exhibited comparable OA severity to joints receiving DMM surgery (both WT and cKO male mice) (**Fig. 4C**). DMM joints of male cKO mice also had increased tibial subchondral bone mineralization density versus non-injured cKO joints (**Fig. 4D**). No significant difference in OA severity was observed among female mice (**Fig. 4B&C, Supp. Fig. 6**). DMM joints of male WT mice exhibited significantly higher synovitis compared to non-injured WT joints (**Supp. Fig. 7**). Additionally, cKO non-injured mice had a trend of increased synovitis, comparable to mice with DMM surgery (**Supp. Fig. 7**). We next performed IHC of PRDM16 to examine whether OA development affects temporal expression of PRDM16 in C57BL/6J mice. At 7 days post-DMM surgery, high expression of PRDM16 was detected both in the articular cartilage and medial meniscus of non-surgery control and injured joints (**Supp. Fig. 8A&B**). Additionally, no signs of OA were detected at this time point. However, by 12-wks post-surgery, PRDM16 expression was significantly decreased in the articular cartilage of both the tibia and femur in the DMM joint but not in the non-injury control (**Supp. Fig. 8A&B**). Moreover, analysis of publicly available bulk RNA-seq datasets from human healthy and OA knee cartilage^44^ revealed decreased PRDM16 gene expression in OA cartilage compared with healthy tissue (**Supp. Fig. 8C**).

**Fig. 4:**
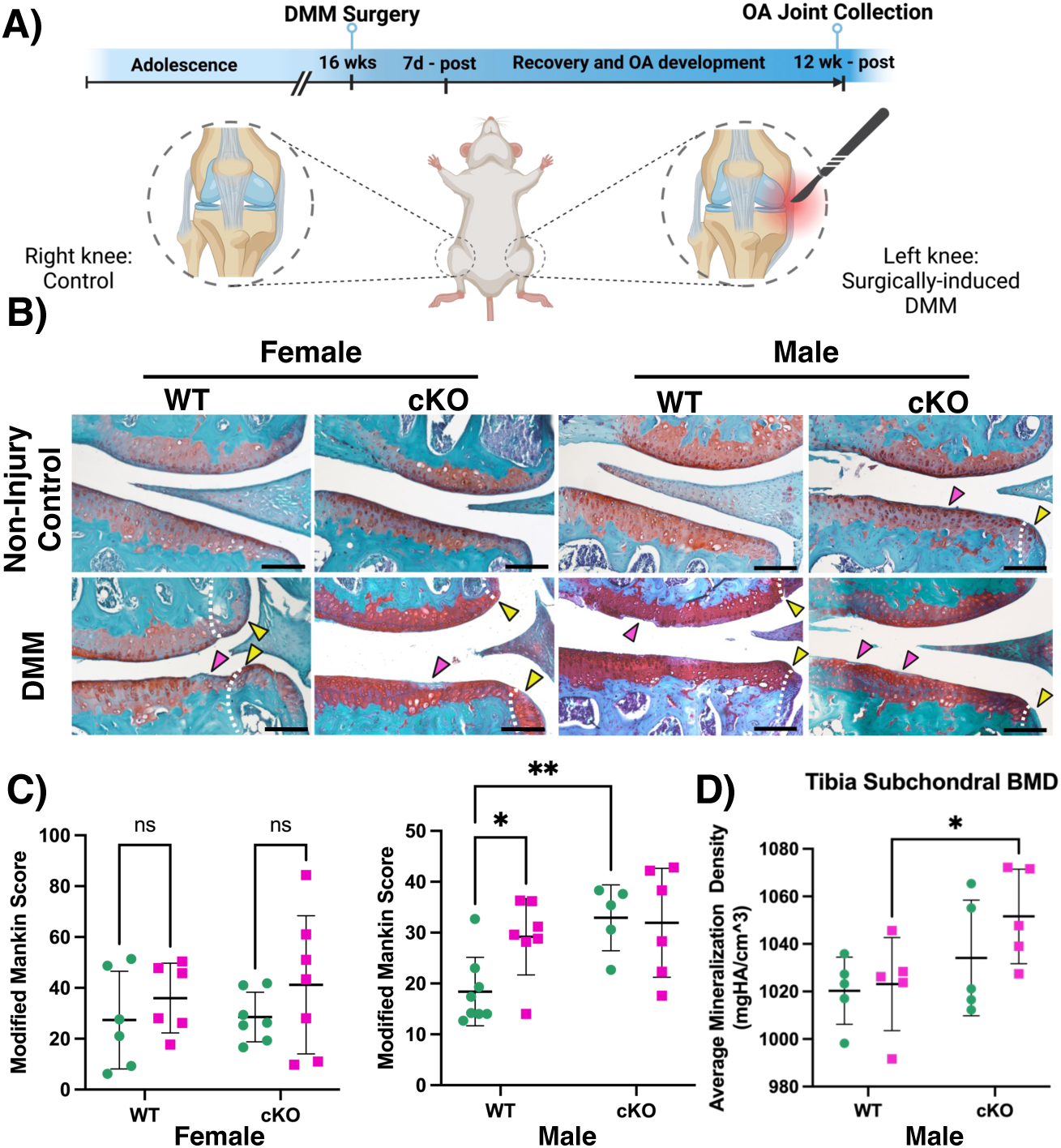
PRDM16 plays an important role in knee joint homeostasis. (**A**) Schematic of DMM surgery and timeline. (**B**) Saf-O/Fast Green staining of DMM and non-surgery control joints from female and male mice (medial side shown). Arrowheads indicate cleft or loss of cartilage (pink) and osteophyte (yellow). Black scale bar is 100µm. (**C**) Modified Mankin grading of non-injury control (turquoise) and DMM (pink) joints. (**D**) Tibia subchondral BMD of non-injury control (turquoise) and DMM (pink) male mice. Two-Way ANOVA with Tukey’s post-hoc (*p < 0.05, **p < 0.01).

### Both over- and knockdown expression of PRDM16 in hiPSC-derived chondrocytes results in abnormal chondrogenesis

To elucidate the functional role of PRDM16 in chondrogenesis, we generated hiPSC lines capable of inducing either OE or KD expression of PRDM16 with doxycycline treatment. The hiPSC chondrogenesis was then carried out according to our previously established protocol^24^, and OE or KD of PRDM16 was induced at the chondroprogenitor (C_P_) stage^3^. Successful induction of desired PRDM16 expression was confirmed with western blot analysis (**Fig. 5A**). Both control and OE hiPSC-derived chondrogenic pellets had significantly higher GAG concentration compared to KD cells (**Fig. 5B&C**). Interestingly, both PRDM16 OE and KD chondrocytes had reduced DNA content as compared to the control pellets (**Fig. 5B&C**). Unsupervised clustering of integrated scRNA-seq datasets of hiPSC-derived chondrocytes from all treatment groups revealed 11 distinct cell populations (**Fig. 5D & Supp. Fig. 9**). Marker genes of each cluster were selected based on unique (+) or increased relative expression (high) of the select gene in its distinct cluster. Only Clusters 0, 1, 2, 3, 5, 6, and 10 from Fig. 5D were further subset and re-clustered for downstream analyses, focusing on the role of PRDM16 in chondrocyte phenotype specification. This process resulted in 5 distinct unique chondrocyte subpopulations (**Fig. 5E-G**). Note that UCMA^+^/ACAN^high^ chondrocytes (Cluster 2) exhibit the highest expression of mature chondrocyte markers including *ACAN*, *EPYC*, and *COL2A1*. KD cells exhibited an increased percentage of SOX5^high^/SOX6^high^ chondrocytes (Cluster 3, blue) compared to the other two treatment groups (KD: 13.5% vs. OE: 4.8%, Control: 7.0%) (**Fig. 5F, Supp. Table 3**). OE chondrocytes exhibited a high percentage of IGFBP5^high^/COL3A1^high^ chondrocytes (Cluster 1, olive) (OE: 47.2% vs. KD: 26.3%, Control: 33.4%) (**Fig. 5F, Supp. Table 3**).

**Fig. 5:**
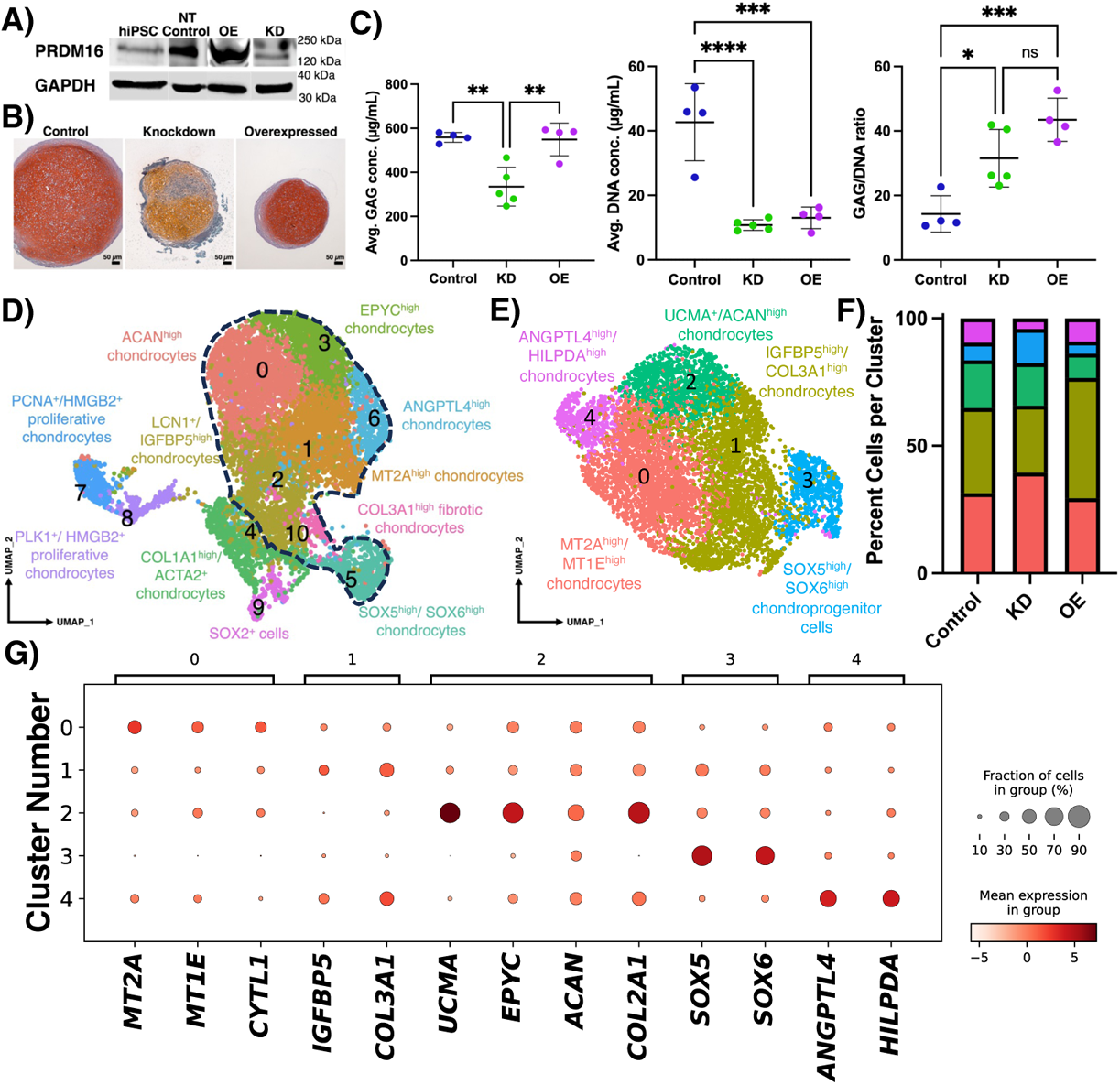
Unsupervised clustering of hiPSC-derived chondrocytes with Control, KD, or OE expression. (**A**) Western blot of hiPSC-derived chondrocytes. (**B**) Saf-O/Fast Green staining of chondrogenic pellets. (**C**) Biochemical analysis of pellet GAG and DNA concentrations. (**D**) scRNA-seq integrated data revealed 11 unique populations conserved among all treatment groups. Black dashed line highlights clusters selected for further subsetting and reclustering. (**E**) Supervised clustering of subset populations reveals 5 distinct chondrocyte populations. (**F**) Percentage of each cluster per treatment group in the subset data. (**G**) Mean expression of select marker gene across subset populations. One-Way ANOVA with Tukey’s post-hoc (*p < 0.05, **p < 0.01, ***p < 0.001, ****p < 0.0001).

### PRDM16 directs cell fate decisions of SOX5^high/^SOX6^high^ chondroprogenitors

By integrating the results from multiple cell differentiation analyses including RNA velocity, Random walk, Palantir, CellRank 2, and Monocle 3, we identified that SOX5^high^/SOX6^high^ chondroprogenitors differentiate into either 1) UCMA^+^ or 2) ANGPTL4^high^ chondrocytes in Control cells (**Fig. 6A and Supp. Fig. 10**). Additionally, Control SOX5^high^/SOX6^high^ chondroprogenitors distribute relatively homogenously across the differentiation trajectory (Palantir pseudotime) (**Fig. 6B**). Fate probability analysis further revealed that Control SOX5^high^/SOX6^high^ chondroprogenitors have a higher fate probability toward ANGPTL4^high^ chondrocytes (67%) vs. UCMA^+^ chondrocytes (33%) (**Fig. 6C and Supp. Fig. 11**). In KD cells, SOX5^high^/SOX6^high^ chondroprogenitors differentiate into either 1) UCMA^+^ or 2) MT2A^high^ chondrocytes (**Fig. 6D**). Interestingly, many of the KD SOX5^high^/SOX6^high^ chondroprogenitors remain undifferentiated (**Fig. 6E**). However, if these cells do successfully differentiate, KD SOX5^high^/SOX6^high^ chondroprogenitors most likely commit towards MT2A^high^ chondrocytes (93%) vs. UCMA^+^ chondrocytes (3%) (**Fig. 6F**). GO functional analysis of DEGs of SOX5^high^/SOX6^high^ chondroprogenitors under KD versus Control conditions revealed that PRDM16 KD was associated with up-regulation of GO terms related to Serine/threonine kinase signaling, axonogenesis, and the TGF-β signaling pathway, and down-regulation of GO terms related to response to hypoxia, limb development, and collagen fibril organization (**Supp. Fig. 12**). In addition, GSEA identified enrichment of the LIM_MAMMARY_STEM_CELL_UP and BOQUEST_STEM_CELL_UP gene sets in the SOX5^high^/SOX6^high^ chondroprogenitor population under KD conditions (**Supp. Fig. 12**). OE SOX5^high^/SOX6^high^ chondroprogenitors differentiate into either 1) IGFBP5^high^, 2) MT2A^high^, or 3) UCMA^+^ chondrocytes, with MT2A^high^ chondrocytes (55%) having the highest fate probability (**Fig. 6G**). While most OE SOX5^high^/SOX6^high^ chondroprogenitors reach terminal states of differentiation, (e.g., UCMA^+^ or MT2A^high^ chondrocytes), a significant number commit to IGFBP5^high^ chondrocytes (24%; **Fig. 6H-I**). GO functional analysis of DEGs in the SOX5^high^/SOX6^high^ chondroprogenitors between OE compared to Control revealed that OE PRDM16 is associated with up-regulation of terms related to actin filament organization, muscle differentiation, and Serine/threonine kinase signaling, and down-regulation of terms related to response to ribonucleoprotein complex biogenesis, ossification, and cartilage development (**Supp. Fig. 13**).

**Fig. 6:**
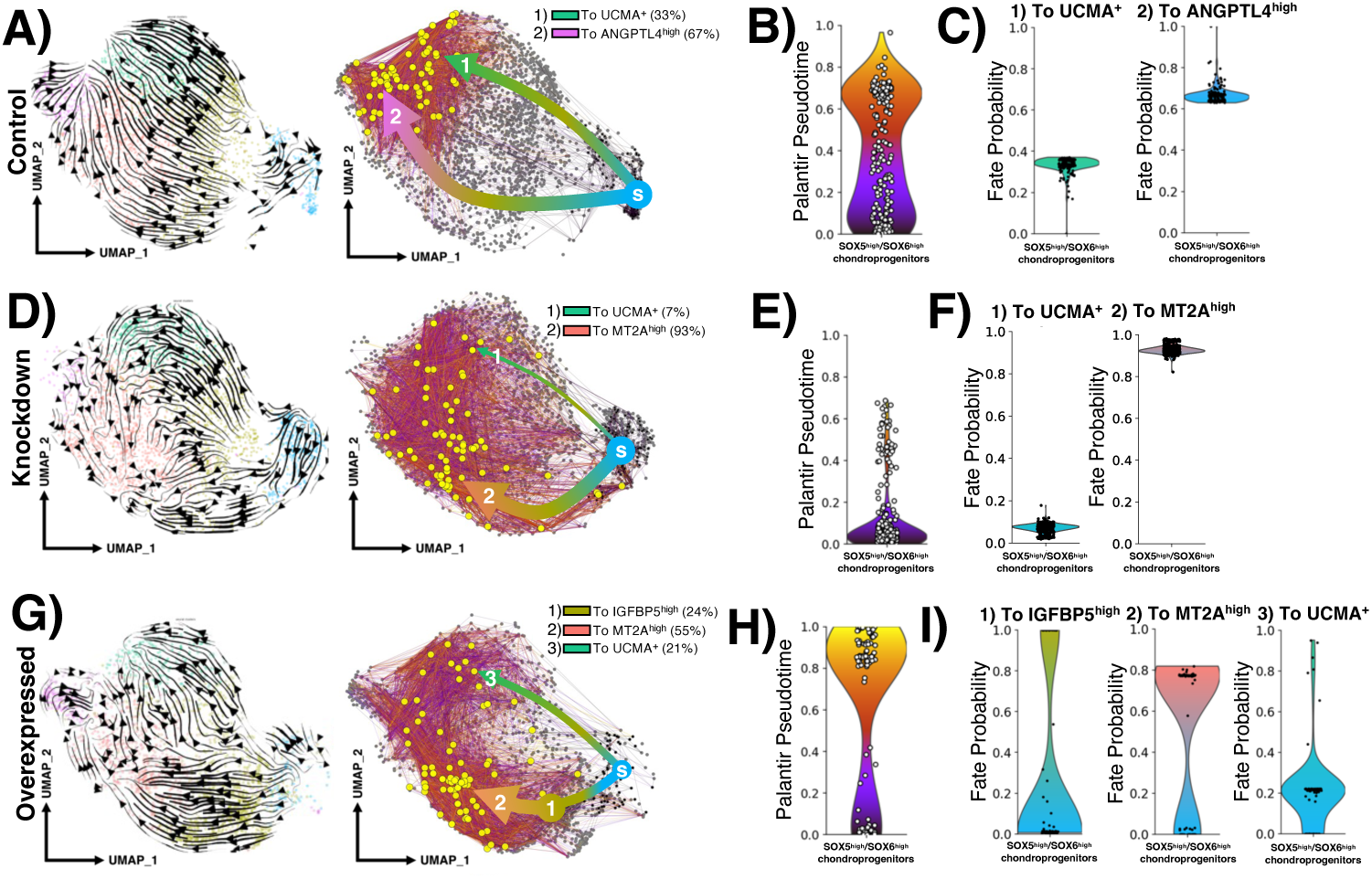
Palantir pseudotime analysis predicts a PRDM16-dependent cell fate determination of SOX5^high^/SOX6^high^ chondroprogenitors. **(A)** Control SOX5^high^/SOX6^high^ chondroprogenitors may differentiate into either 1) UCMA^+^ or 2) ANGPTL4^high^ chondrocytes based on RNA velocity (left) and random walk (right) analyses (s = start of random walk, yellow dots = end of random walk; numbers indicate average fate probability to a given lineage determined by CellRank). Thicker arrow lengths indicate higher fate probability. **(B)** Control SOX5^high^/SOX6^high^ chondroprogenitor exhibited a relatively homogenous distributions of cells across the differentiation trajectory. **(C)** Fate probability analysis revealed that Control SOX5^high^/SOX6^high^ chondroprogenitors exhibit increased fate probability toward ANGPTL4^high^ vs. UCMA^+^ chondrocytes. **(D)** KD SOX5^high^/SOX6^high^ chondroprogenitors may differentiate into either 1) UCMA^+^ or 2) MT2A^high^ chondrocytes. **(E)** Many KD SOX5^high^/SOX6^high^ chondroprogenitors remained in the undifferentiated stage, however **(F)** if successfully differentiated, KD SOX5^high^/SOX6^high^ likely commit towards the MT2A^high^ chondrocyte fate rather towards UCMA^+^ chondrocytes. **(G)** OE SOX5^high^/SOX6^high^ chondroprogenitors may differentiate into 1) IGFBP5^high^, 2) MT2A^high^, or 3) UCMA^+^ chondrocytes, with the highest fate probability being towards 2) MT2A^high^ chondrocytes**. (H)** Most OE SOX5^high^/SOX6^high^ chondroprogenitors reach terminal states of differentiation**. (I)** OE SOX5^high^/SOX6^high^ chondroprogenitors exhibit a 24% fate commitment towards IGFBP5^high^ chondrocytes.

### PRDM16 regulates genes involved in chondrogenesis by altering DNA binding and/or chromatin accessibility

To further elucidate the genetic and epigenetic mechanisms by which PRDM16 regulates chondrogenesis, we performed CUT&RUN-seq using antibodies against PRDM16, H3K4me3, and H3K9me1 on hiPSC-derived chondrogenic pellets with Control or KO expression of PRDM16. Coverage heatmaps revealed that peaks for PRDM16 DNA binding sites and histone marks were significantly reduced in PRDM16 KO (**Fig. 7A**). Interestingly, PRDM16 is highly enriched in promoter/enhancer regions of select DNA binding sites and H3K4me3 ( >30%) but not H3K9me1 (∼1%) (**Fig. 7B**). Next, we integrated our scRNA-seq and CUT&RUN-seq datasets to determine if genes that had differential expression patterns between WT and KO were mediated via PRDM16 DNA binding and/or histone modification at promoter/enhancer regions. For example, along the differentiation trajectory from SOX5^high^/SOX6^high^ chondroprogenitors to UCMA^+^ chondrocytes, we identified several genes including *SMOC2*, *ENPP1*, *ADAMTS19*, *HAND2*, and *HGF* that had opposite expression trends between Control and KD pellets, suggesting that these genes may be regulated by PRDM16 through promoter/enhancer binding and/or H3K4me3 modification (**Fig. 7C**). Specificially, these genes were up-regulated in Control but down-regulated in PRDM16 KD pellets across this differentiation trajectory. Consistent with the transcriptomic data, KO pellets had significantly reduced enrichment peaks at the promoter regions of *SMOC2* (likely mediated by DNA binding) and *HAND2* (likely mediated by H3K4me3 modification) (**Fig. 7D&E**). In the OE pellets, several genes including *MEF2C*, *FADS2*, and *MRNIP* were up-regulated along the differentiation trajectory of SOX5^high^/SOX6^high^ chondroprogenitors to IGFBP5^high^ chondrocytes, correlating with increased PRDM16 abundance (**Fig. 7F-H**).

**Fig. 7:**
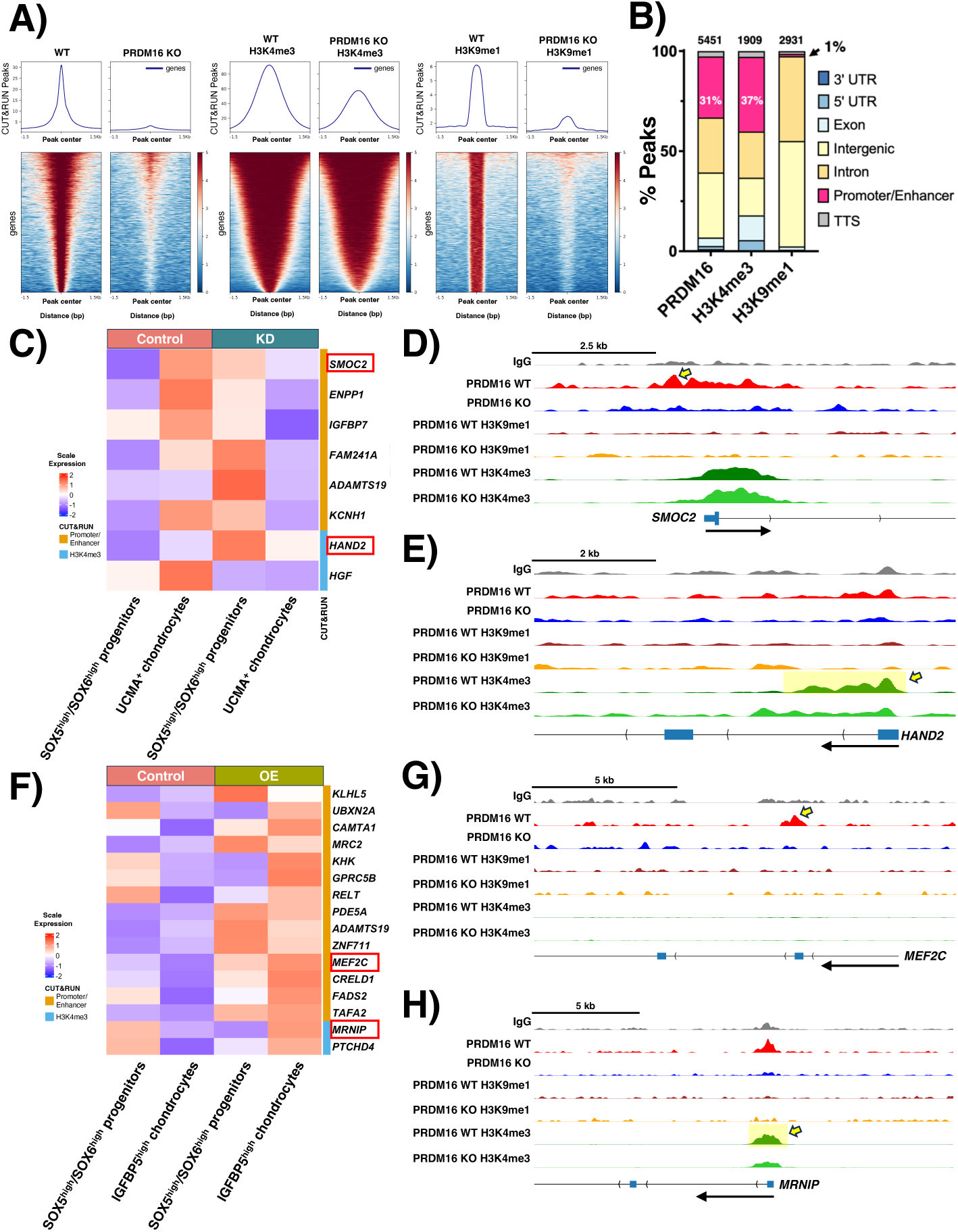
KD and OE PRDM16 alter chondrogenic gene expression by altering DNA binding and chromatin states. CUT&RUN sequencing targeting PRDM16, H3K4me3, and H3K9me1 were performed on WT (Control) and PRDM16 KO chondrogenic pellets. (**A**) Coverage heatmaps showing ± 1.5 kb from peak centers of PRDM16 DNA binding, H3K4me3, and H3K9me1 in WT and KO cells. A significant reduction in peaks was observed in PRDM16 KO samples. (**B**) Genomic annotation of enriched peaks in WT cells demonstrates preferential enrichment at promoter/enhancer regions for DNA binding and H3K4me3 marks. (**C**) Heatmap illustrating genes with opposite expression patterns between Control and PRDM16 KD cells along the differentiation trajectory of SOX5^high^/SOX6^high^ chondroprogenitors to UCMA^+^ chondrocytes. Genes annotated with orange or blue were also identified by CUT&RUN-seq as associated with DNA binding or H3K4me3 modification, respectively. Representative genome track plots for (**D**) *SMOC2* (PRDM16 DNA binding) and **(E)** *HAND2* (PRDM16 H3K4me3 modification). Yellow arrows indicate peak location. (**F**) Heatmap depicting genes with opposite expression patterns between Control and PRDM16 OE pellets along the differentiation trajectory of SOX5^high^/SOX6^high^ chondroprogenitors to IGFBP5^high^ chondrocytes. Representative genome track plots for (**G**) *MEF2C* (PRDM16 DNA binding) and (**H**) *MRNIP* (PRDM16 H3K4me3 modification).

Next, to identify chondrogenic markers that are potentially regulated by PRDM16, we curated genes from cartilage-associated GO terms (**Supp. Table 4**) and intersected them with genes identified from the CUT&RUN-seq. We found 10 cartilage-associated genes whose promoter/enhancer regions were enriched for PRDM16 DNA binding, and 3 cartilage-associated genes whose promoter/enhancer regions were enriched for H3K4me3 modification. No genes exhibited concurrent enrichment for both PRDM16 binding and H3K4me3/H3K9me1(**Fig. 8A**). Among the 10 cartilage-associated genes enriched for PRDM16 DNA binding, *MEF2C*, *ITGA2*, *ARID5A*, and *TRIP11* showed expression patterns positively associated with PRDM16 expression, while *RUNX1* had a negative association with PRDM16 expression (**Fig. 8B**). Interestingly, *SOX9* expression was decreased with KD PRDM16, but the lowest expression levels were observed in cells with OE PRDM16. Expression of *MEF2C* and *TRIP11* were decreased in the KD cells, and up-regulated in OE cells across all chondrogenic subpopulations (**Fig. 8C**). Expression of *ITGA2* was low across all populations. The proposed mechanism by which PRDM16 regulates genes associated with chondrogenesis and genes associated with chondrocyte hypertrophy has been illustrated schematically (**Fig. 8D**).

**Fig. 8:**
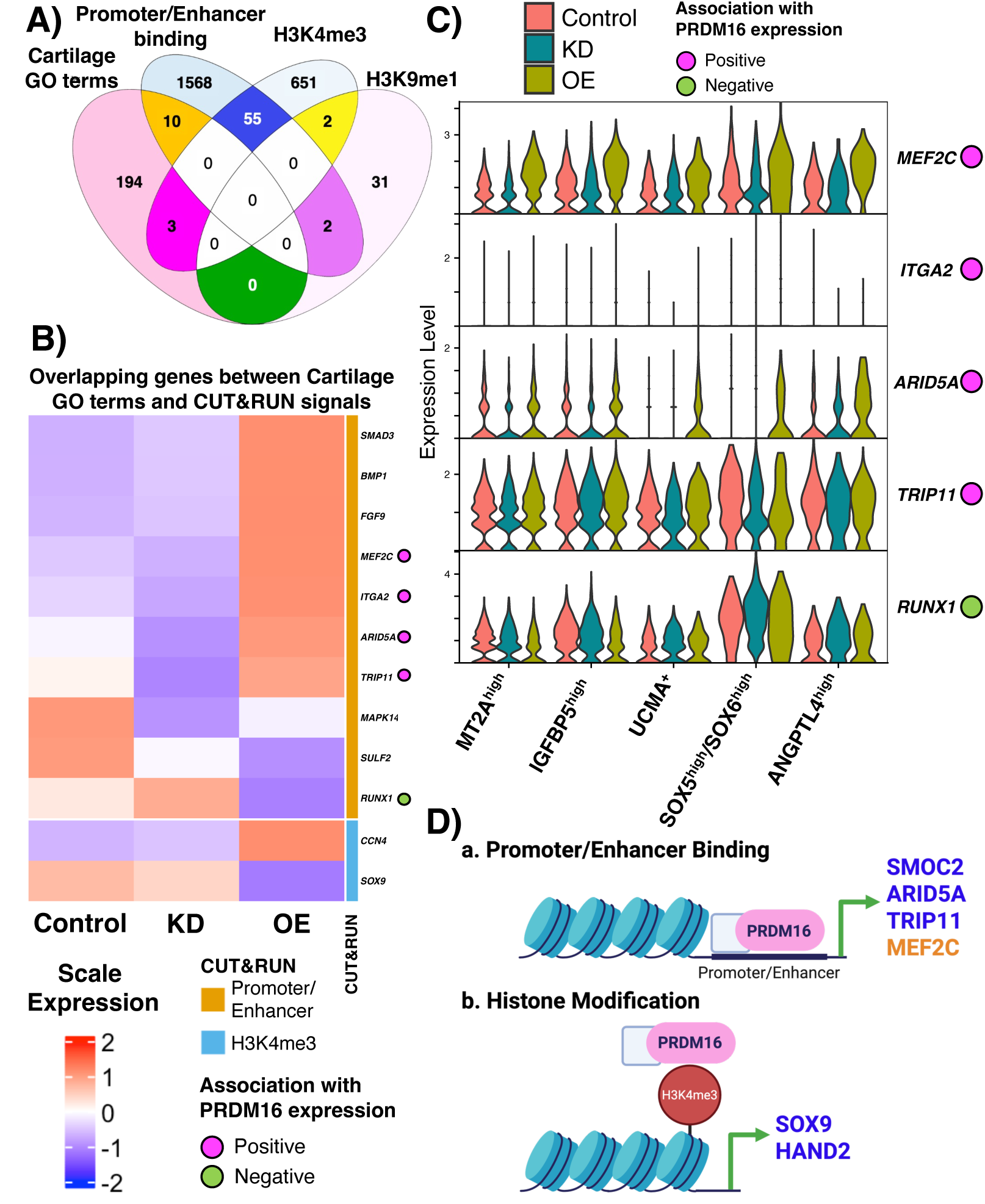
PRDM16 regulates chondrogenic and hypertrophic genes via promoter/enhancer binding and histone modification. (**A**) Venn diagram of overlap between genes associated with cartilage-related GO terms, PRDM16 DNA binding, and histone modifications. (**B**) Heatmap and (**C**) violin plots showing expression levels of genes overlapping between cartilage GO terms and CUT&RUN signals across different PRDM16 conditions or chondrocyte populations, respectively. Genes including *MEF2C*, *ITGA2*, *ARID5A,* and *TRIP11* had a positive association with PRDM16 (pink dot), while RUNX1 had a negative association with PRDM16 expression (green dot). (**D**) Schematic of the potential mechanism by which PRDM16 regulates genes associated with chondrogenesis (blue) and genes associated with chondrocyte hypertrophy (orange).

## Discussion

The findings of this study demonstrate that PRDM16 acts as a transcriptional and epigenetic regulator of chondrogenesis and articular chondrocyte phenotypes in knee cartilage. We identified that PRDM16 regulates expression of *MEF2C*, *ARID5A*, and *TRIP11* through binding of promoter or enhancer regions. Furthermore, we elucidated that PRDM16 modulates the H3K4me3 activation mark on several essential TFs, including *HAND2* and *SOX9*, required for normal chondrogenesis. These findings align with previous work that shows loss of PRDM16 results in craniofacial cartilage defects in zebrafish driven by reduced H3K4me3 (gene activation)^10^. Interestingly, in *Sox2-Cre^+/Tg^;Prdm16^flox/+^* mice, loss of PRDM16 is linked to a significant decrease in H3K9 but not H3K4 methylation in the palatal shelves, although the role of PRDM16 in knee cartilage development has not been previously investigated^10^.

PRDM16 expression is detectable in the mouse limb bud mesenchyme as early as E10.5^45^, suggesting it may play a role in knee morphogenesis. However, using our *Col2a1Cre;Prdm16^flox/flox^* model, we observed no significant differences in percent length of chondrocyte zonal structures between WT and cKO mice at E15.5. However, by E18.5, loss of Prdm16 led to an accumulation of resting chondrocytes and abbreviated proliferative and hypertrophic zones. Furthermore, E18.5 cKO mice exhibited increased MKI67 expression in their columnar proliferative chondrocytes, indicating delayed development through differentiation. cKO mice also exhibited markedly reduced MKI67 expression in the perichondrium indicating reduced proliferation in the knee joint compartment. The perichondrium negatively regulates chondrocyte maturation and hypertrophy, and its loss is associated with an extended zone of immature chondrocytes^46^. Thus, reduced perichondrial proliferation in cKO mice may explain the decrease in columnar proliferative and hypertrophic chondrocytes observed at E18.5. Collectively, these data support a critical role for PRDM16 in regulating the transition from resting to proliferative chondrocytes, and in coordinating proper long bone development.

As mice with gKO of Prdm16 are neonatal lethal^45^, little is known as to whether PRDM16 is necessary for maintaining postnatal joint homeostasis. Using our cKO model, we examined cartilage development of adolescent (4-wk-old) and adult (12-wk-old) mice. We found that 4-week-old Prdm16 cKO mice exhibited a trend of decreased body weight versus WT, in line with a previous study reporting that Prdm16^+/-^ mice have mildly reduced body mass than WT mice^13^. Although sex was investigated independently, no significant differences were observed between female and male cKO mice during development. Kaneda-Nakashima et al. have reported that one-year-old Prdm16^+/-^ mice display disrupted organization of the growth plate^13^, but we did not observe any apparent differences in the knee epiphysial cartilage between WT and cKO mice at either 4- or 12-weeks. This difference may reflect the extent and cellular distribution of Prdm16 loss, as our model targets osteochondral lineage cells, whereas the prior study involved a global heterozygous deletion.

Our Prdm16 cKO mice also exhibited impaired bone development in the hind limbs at the early stages of development. Interestingly, both female and male 4-week-old cKO mice had no ossification of the medial meniscus and reduced ossification of the lateral meniscus compared to WT. To the best of our knowledge, delayed meniscal ossification resulting from loss of PRDM16 has not been previously reported, despite this developmental process normally being initiated by 2 weeks and readily detectable by 4 weeks of age in rodents^19^. Although decreased BV/TV of several joint bones (e.g., tibial tuberosity, patella) in cKO mice mostly recovered by 12-weeks of age, cKO mice still exhibited reduced medial meniscal ossification. The contribution of PRDM16 on bone formation is still poorly understood and may vary depending on the ossification process (intramembranous versus endochondral). For example, loss of *Prdm16* in Meckel’s cartilage leads to defective chondrocyte maturation and mandibular hypoplasia, indicating a requirement of PRDM16 for endochondral ossification^47^. Furthermore, PRDM16 has been reported to negatively regulate terminal osteogenic differentiation during intramembranous ossification. Because osteoblast progenitors arise from multiple developmental lineages—including chondrocytes, perichondrial cells, periosteal cells, and marrow stromal cells – lineage-specific Col2a1 and Prdm16 expression among these lineages may explain the variable recovery rates observed across bone compartments in Prdm16 cKO mice^48^. Overall, our findings demonstrate that loss of Prdm16 in osteochondral lineage cells delays, but does not fully inhibit, bone formation in the knee joint.

To investigate the role of PRDM16 in articular cartilage homeostasis and response to injury, we performed DMM surgery to induce OA in WT and cKO mice. Increased cartilage degradation, osteophyte formation, and synovitis were observed in the DMM joints as compared to non-surgery joints in WT male mice, but not in WT female mice. This is consistent with previous findings that mouse OA severity is markedly higher in males than females after DMM^49, 50^. Interestingly, cKO male mice had comparable OA severity between DMM and non-surgery joints, suggesting that PRDM16 may exert a chondroprotective function. This proposition is further supported by the significant reduction of PRDM16 expression observed in the cartilage of mouse DMM joints, and in human osteoarthritic cartilage^44^.

In order to further identify the molecular mechanisms by which PRDM16 regulates chondrogenesis, we integrated scRNA-seq and CUT&RUN-seq datasets from our hiPSC model with modulated PRDM16 expression. Interestingly, both PRDM16 KD and OE exhibited reduced DNA content versus Control, implying a link between PRDM16 and chondrocyte viability. This finding is consistent with previous reports that show PRDM16 is involved in stem cell maintenance by regulating oxidative stress^51, 52^. PRDM16 KD chondrocytes also exhibited significantly decreased Saf-O staining and reduced GAG concentration, while OE expressed comparable GAG production to Control. These results suggest that PRDM16 is essential for promoting chondrogenesis and that while OE cells have poor viability, surviving cells retain the capacity to undergo normal chondrogenic differentiation.

Our multiomic analyses revealed that PRDM16 modulation alters chromatin accessibility and DNA binding of genes essential for chondrogenesis, leading to distinct distributions of chondrocyte fates. KD cells displayed an accumulation of SOX5^high^/SOX6^high^ chondroprogenitors, consistent with GSEA, indicating arrested differentiation. In other types of stem cells, including hematopoietic stem cells and neural stem cells, loss of PRDM16 reduces stemness^53, 54^. PRDM16 is also required for brown adipogenesis, and depletion thereof leads to failure of brown fat differentiation, leaving cells in a preadipocyte-like state^55^. Similarly, our results suggest that loss of PRDM16 causes chondroprogenitors to become arrested in an undifferentiated state, reflecting impaired chondrogenic differentiation. Moreover, dysregulated expression of essential genes for cartilage and bone development including *SMOC2*, *HAND2*, *ENPP1*^56^ (**Fig. 7C**), *SOX9*, *MEF2C*, *ARID5A*, and *TRIP11* (**Fig. 8B&C**) further explains the impaired chondrogenesis observed in PRDM16 KD pellets. For example, SMOC2 is known to negatively regulate chondrogenic differentiation, and its loss leads to abnormal limb development^57^. Furthermore, HAND2 represses early chondrogenesis while later slowing chondrocyte maturation^58^. Elevated *SMOC2* and *HAND2* expression in *SOX5^high^/SOX6^high^* chondroprogenitors likely contributes to the persistence of undifferentiated precursors in KD conditions. MEF2C is of particular interest as its expression decreases in KD but is up-regulated in OE cells across all chondrocyte subpopulations. MEF2C is an essential TF required for chondrocyte hypertrophy and endochondral ossification^59^, providing a mechanistic explanation for the impaired bone mineralization observed in Prdm16 cKO mice. Importantly, we identified a putative PRDM16 binding site within a validated enhancer of MEF2C (ENCODE Accession: EH38E2389409), suggesting that PRDM16 plays a direct regulatory role in this mechanism. Notably, several CUT&RUN-identified genes (presumably regulated by PRDM16) did not consistently mirror the PRDM16 expression patterns observed in chondrocyte subpopulations. It is likely that our bulk CUT&RUN-seq data lacks the resolution to fully reflect PRDM16-dependent gene expression changes across the chondrocyte subpopulations identified by scRNA-seq. However, expression patterns of numerous CUT&RUN-identified genes are opposite between Control and KD or OE conditions. This implies that PRDM16 exhibits phenotype-specific chondrocyte regulatory functions.

Additionally, OE PRDM16 cells displayed increased IGFBP5^high^ chondrocytes. IGFBP5 stabilizes IGF-1, promoting chondrogenesis^60, 61^, and IGFBP5 is also up-regulated during normal early chondrocyte differentiation^24, 62^. Surprisingly, OE PRDM16 did not enhance chondrogenic differentiation, as evidenced by the reduced DNA content and decreased UCMA^+^ cell population (vs. Control). Our multiomic analysis indicated that SOX9, a master TF driving chondrogenesis, exhibited its lowest expression levels in OE cells. Additionally, RUNX1, known to cooperate with the SOX-trio in promoting chondrocyte differentiation, displayed an inverted expression pattern relative to PRDM16^63^. Moreover, elevated MEF2C in OE cells may further drive chondrocyte hypertrophy. Together these findings suggest that both insufficient and excessive PRDM16 expressions disrupt the complex transcriptional balance required for normal chondrogenesis.

Several future directions warrant exploration. First, inducible Cre lines (e.g., Acan-CreER^T2^) would allow temporal dissection of PRDM16 function in cartilage homeostasis, injury response, and aging. Additionally, treating hiPSC-derived chondrocytes with inflammatory cytokines, such as IL-1, may provide clarity on how PRDM16 is regulated under catabolic conditions, providing mechanistic insight into the downregulation of PRDM16 in OA. Furthermore, single-cell CUT&RUN approaches may resolve the dynamic, cell state-specific, epigenetic functions of PRDM16 during chondrogenesis and endochondral ossification.

Overall, we have demonstrated that PRDM16 deficiency delays ossification of multiple joint structures and increases susceptibility to OA. Furthermore, through promoter/enhancer binding and regulation of H3K4me3 deposition, PRDM16 governs the expression of key chondrogenic regulators including SOX9, ARID5A, SMOC2, HAND2, and hypertrophic driver MEF2C. Together, our findings establish PRDM16 as an essential genetic and epigenetic regulator of chondrogenesis and chondrocyte phenotype specification in the knee joint.

## Acknowledgments

The authors thank UR Genomics Research Center for assisting with scRNA-seq. This study was supported in part by NIH grants AR075899, AR082403, UR and C-COMP P30 pilots, Orthopaedic Research and Education Foundation, Arthritis National Research Foundation, and NSF GRFP.

## References

1. Hunter DJ, March L, Chew M. Osteoarthritis in 2020 and beyond: a Lancet Commission. Lancet. 2020;396(10264):1711–2. Epub 20201104. doi: 10.1016/s0140-6736(20)32230-3. PubMed PMID: 33159851.

2. Lee JS, Shim DW, Kang KY, Chae DS, Lee WS. Method Categorization of Stem Cell Therapy for Degenerative Osteoarthritis of the Knee: A Review. Int J Mol Sci. 2021;22(24). Epub 20211211. doi: 10.3390/ijms222413323. PubMed PMID: 34948119; PMCID: PMC8704290.

3. Wu C-L, Dicks A, Steward N, Tang R, Katz DB, Choi Y-R, Guilak F. Single cell transcriptomic analysis of human pluripotent stem cell chondrogenesis. Nature Communications. 2021;12(1). doi: 10.1038/s41467-020-20598-y.

4. Yamato G, Kawai T, Shiba N, Ikeda J, Hara Y, Ohki K, Tsujimoto SI, Kaburagi T, Yoshida K, Shiraishi Y, Miyano S, Kiyokawa N, Tomizawa D, Shimada A, Sotomatsu M, Arakawa H, Adachi S, Taga T, Horibe K, Ogawa S, Hata K, Hayashi Y. Genome-wide DNA methylation analysis in pediatric acute myeloid leukemia. Blood Adv. 2022;6(11):3207–19. doi: 10.1182/bloodadvances.2021005381. PubMed PMID: 35008106; PMCID: PMC9198913.

5. Kajimura S, Seale P, Kubota K, Lunsford E, Frangioni JV, Gygi SP, Spiegelman BM. Initiation of myoblast to brown fat switch by a PRDM16-C/EBP-beta transcriptional complex. Nature. 2009;460(7259):1154–8. Epub 20090729. doi: 10.1038/nature08262. PubMed PMID: 19641492; PMCID: PMC2754867.

6. Li J, Godoy MI, Lu Y, Zhang AJ, Diamante G, Rathbun E, Tian M, Ahn IS, Cebrian-Silla A, Alvarez-Buylla A, Yang X, Novitch BG, Carmichael ST, Zhang Y. Prdm16 regulates the postnatal fate of embryonic radial glia via Vcam1-dependent mechanisms. Nat Commun. 2025;16(1):6659. Epub 20250719. doi: 10.1038/s41467-025-60895-y. PubMed PMID: 40683857; PMCID: PMC12276310.

7. Van Wauwe J, Kemps H, Vrancaert P, Mahy A, Schellingen R, Grootaert MOJ, Beerens M, Luttun A. PRDM16, a new kid on the block in cardiovascular health and disease. Cardiovasc Res. 2025;121(8):1156–72. doi: 10.1093/cvr/cvaf089. PubMed PMID: 40439125; PMCID: PMC12310283.

8. Jordan VK, Zaveri HP, Scott DA. 1p36 deletion syndrome: an update. Appl Clin Genet. 2015;8:189–200. Epub 20150827. doi: 10.2147/tacg.S65698. PubMed PMID: 26345236; PMCID: PMC4555966.

9. Motch Perrine SM, Wu M, Holmes G, Bjork BC, Jabs EW, Richtsmeier JT. Phenotypes, Developmental Basis, and Genetics of Pierre Robin Complex. J Dev Biol. 2020;8(4). Epub 20201205. doi: 10.3390/jdb8040030. PubMed PMID: 33291480; PMCID: PMC7768358.

10. Shull LC, Sen R, Menzel J, Goyama S, Kurokawa M, Artinger KB. The conserved and divergent roles of Prdm3 and Prdm16 in zebrafish and mouse craniofacial development. Dev Biol. 2020;461(2):132–44. Epub 20200208. doi: 10.1016/j.ydbio.2020.02.006. PubMed PMID: 32044379; PMCID: PMC7198358.

11. Jugessur A, Shi M, Gjessing HK, Lie RT, Wilcox AJ, Weinberg CR, Christensen K, Boyles AL, Daack-Hirsch S, Nguyen TT, Christiansen L, Lidral AC, Murray JC. Maternal genes and facial clefts in offspring: a comprehensive search for genetic associations in two population-based cleft studies from Scandinavia. PLoS One. 2010;5(7):e11493. Epub 20100709. doi: 10.1371/journal.pone.0011493. PubMed PMID: 20634891; PMCID: PMC2901336.

12. Bjork BC, Turbe-Doan A, Prysak M, Herron BJ, Beier DR. Prdm16 is required for normal palatogenesis in mice. Hum Mol Genet. 2010;19(5):774–89. Epub 20091211. doi: 10.1093/hmg/ddp543. PubMed PMID: 20007998; PMCID: PMC2816611.

13. Kaneda-Nakashima K, Igawa K, Suwanruengsri M, Naoyuki F, Ichikawa T, Funamoto T, Kurogi S, Sekimoto T, Yamashita Y, Chosa E, Yamaguchi R, Morishita K. Role of Mel1/Prdm16 in bone differentiation and morphology. Exp Cell Res. 2022;410(2):112969. Epub 20211207. doi: 10.1016/j.yexcr.2021.112969. PubMed PMID: 34883111.

14. Chou CH, Wu CC, Song IW, Chuang HP, Lu LS, Chang JH, Kuo SY, Lee CH, Wu JY, Chen YT, Kraus VB, Lee MT. Genome-wide expression profiles of subchondral bone in osteoarthritis. Arthritis Res Ther. 2013;15(6):R190. doi: 10.1186/ar4380. PubMed PMID: 24229462; PMCID: PMC3979015.

15. Harasymowicz NS, Choi YR, Wu CL, Iannucci L, Tang R, Guilak F. Intergenerational Transmission of Diet-Induced Obesity, Metabolic Imbalance, and Osteoarthritis in Mice. Arthritis Rheumatol. 2020;72(4):632–44. Epub 20200305. doi: 10.1002/art.41147. PubMed PMID: 31646754; PMCID: PMC7113102.

16. O’Conor CJ, Ramalingam S, Zelenski NA, Benefield HC, Rigo I, Little D, Wu CL, Chen D, Liedtke W, McNulty AL, Guilak F. Cartilage-Specific Knockout of the Mechanosensory Ion Channel TRPV4 Decreases Age-Related Osteoarthritis. Sci Rep. 2016;6:29053. Epub 20160708. doi: 10.1038/srep29053. PubMed PMID: 27388701; PMCID: PMC4937413.

17. Moody HR, Heard BJ, Frank CB, Shrive NG, Oloyede AO. Investigating the potential value of individual parameters of histological grading systems in a sheep model of cartilage damage: the Modified Mankin method. J Anat. 2012;221(1):47–54. Epub 20120517. doi: 10.1111/j.1469-7580.2012.01513.x. PubMed PMID: 22591160; PMCID: PMC3390533.

18. Krenn V, Morawietz L, Häupl T, Neidel J, Petersen I, König A. Grading of chronic synovitis--a histopathological grading system for molecular and diagnostic pathology. Pathol Res Pract. 2002;198(5):317–25. doi: 10.1078/0344-0338-5710261. PubMed PMID: 12092767.

19. Gamer LW, Xiang L, Rosen V. Formation and maturation of the murine meniscus. J Orthop Res. 2017;35(8):1683–9. Epub 20161006. doi: 10.1002/jor.23446. PubMed PMID: 27664939.

20. Conn HJ LR. Biological Stains. 9th ed: Waverly Press; 1977.

21. Lillie RD FH. Histopathologic Technic and Practical Histochemistry. 4th ed: McGraw-Hill; 1976.

22. Adkar SS, Wu CL, Willard VP, Dicks A, Ettyreddy A, Steward N, Bhutani N, Gersbach CA, Guilak F. Step-Wise Chondrogenesis of Human Induced Pluripotent Stem Cells and Purification Via a Reporter Allele Generated by CRISPR-Cas9 Genome Editing. Stem Cells. 2019;37(1):65–76. Epub 20181031. doi: 10.1002/stem.2931. PubMed PMID: 30378731; PMCID: PMC6312762.

23. Dicks AR, Steward N, Guilak F, Wu CL. Chondrogenic Differentiation of Human-Induced Pluripotent Stem Cells. Methods Mol Biol. 2023;2598:87–114. doi: 10.1007/978-1-0716-2839-3_8. PubMed PMID: 36355287; PMCID: PMC9830630.

24. Wu C-L, Dicks A, Steward N, Tang R, Katz DB, Choi Y-R, Guilak F. Single cell transcriptomic analysis of human pluripotent stem cell chondrogenesis. Nature Communications. 2021;12(1):362. doi: 10.1038/s41467-020-20598-y.

25. PM Ragan VC, HH Hung, K Masuda, EJ Thonar, EC Arner, AJ Grodzinsky, JD Sandy. Archives of Biochemistry and Biophysics2000. 383 p.

26. Hao Y, Stuart T, Kowalski MH, Choudhary S, Hoffman P, Hartman A, Srivastava A, Molla G, Madad S, Fernandez-Granda C, Satija R. Dictionary learning for integrative, multimodal and scalable single-cell analysis. Nat Biotechnol. 2024;42(2):293–304. Epub 20230525. doi: 10.1038/s41587-023-01767-y. PubMed PMID: 37231261; PMCID: PMC10928517.

27. Hafemeister C, Satija R. Normalization and variance stabilization of single-cell RNA-seq data using regularized negative binomial regression. Genome Biol. 2019;20(1):296. Epub 20191223. doi: 10.1186/s13059-019-1874-1. PubMed PMID: 31870423; PMCID: PMC6927181.

28. Liu Q, He F, Zhou P, Xie M, Wang H, Yang H, Huo W, Zhang M, Yu S, Wang M. HMGB2 promotes chondrocyte proliferation under negative pressure through the phosphorylation of AKT. Biochim Biophys Acta Mol Cell Res. 2021;1868(11):119115. Epub 20210730. doi: 10.1016/j.bbamcr.2021.119115. PubMed PMID: 34333060.

29. Jing Y, Jiang X, Ji Q, Wu Z, Wang W, Liu Z, Guillen-Garcia P, Esteban CR, Reddy P, Horvath S, Li J, Geng L, Hu Q, Wang S, Belmonte JCI, Ren J, Zhang W, Qu J, Liu GH. Genome-wide CRISPR activation screening in senescent cells reveals SOX5 as a driver and therapeutic target of rejuvenation. Cell Stem Cell. 2023;30(11):1452–71.e10. Epub 20231012. doi: 10.1016/j.stem.2023.09.007. PubMed PMID: 37832549.

30. Taniguchi N, Caramés B, Ronfani L, Ulmer U, Komiya S, Bianchi ME, Lotz M. Aging-related loss of the chromatin protein HMGB2 in articular cartilage is linked to reduced cellularity and osteoarthritis. Proc Natl Acad Sci U S A. 2009;106(4):1181–6. Epub 20090112. doi: 10.1073/pnas.0806062106. PubMed PMID: 19139395; PMCID: PMC2633567.

31. McInnes L, Healy J, Melville J. Umap: Uniform manifold approximation and projection for dimension reduction. arXiv preprint arXiv:180203426. 2018.

32. Xu S, Hu E, Cai Y, Xie Z, Luo X, Zhan L, Tang W, Wang Q, Liu B, Wang R, Xie W, Wu T, Xie L, Yu G. Using clusterProfiler to characterize multiomics data. Nat Protoc. 2024;19(11):3292–320. Epub 20240717. doi: 10.1038/s41596-024-01020-z. PubMed PMID: 39019974.

33. Bergen V, Lange M, Peidli S, Wolf FA, Theis FJ. Generalizing RNA velocity to transient cell states through dynamical modeling. Nat Biotechnol. 2020;38(12):1408–14. Epub 20200803. doi: 10.1038/s41587-020-0591-3. PubMed PMID: 32747759.

34. Bray NL, Pimentel H, Melsted P, Pachter L. Near-optimal probabilistic RNA-seq quantification. Nat Biotechnol. 2016;34(5):525–7. Epub 20160404. doi: 10.1038/nbt.3519. PubMed PMID: 27043002.

35. Weiler P, Lange M, Klein M, Pe’er D, Theis F. CellRank 2: unified fate mapping in multiview single-cell data. Nat Methods. 2024;21(7):1196–205. Epub 20240613. doi: 10.1038/s41592-024-02303-9. PubMed PMID: 38871986; PMCID: PMC11239496.

36. Setty M, Kiseliovas V, Levine J, Gayoso A, Mazutis L, Pe’er D. Characterization of cell fate probabilities in single-cell data with Palantir. Nat Biotechnol. 2019;37(4):451–60. Epub 20190321. doi: 10.1038/s41587-019-0068-4. PubMed PMID: 30899105; PMCID: PMC7549125.

37. Cao J, Spielmann M, Qiu X, Huang X, Ibrahim DM, Hill AJ, Zhang F, Mundlos S, Christiansen L, Steemers FJ, Trapnell C, Shendure J. The single-cell transcriptional landscape of mammalian organogenesis. Nature. 2019;566(7745):496–502. Epub 20190220. doi: 10.1038/s41586-019-0969-x. PubMed PMID: 30787437; PMCID: PMC6434952.

38. Qiu X, Mao Q, Tang Y, Wang L, Chawla R, Pliner HA, Trapnell C. Reversed graph embedding resolves complex single-cell trajectories. Nat Methods. 2017;14(10):979–82. Epub 20170821. doi: 10.1038/nmeth.4402. PubMed PMID: 28825705; PMCID: PMC5764547.

39. Trapnell C, Cacchiarelli D, Grimsby J, Pokharel P, Li S, Morse M, Lennon NJ, Livak KJ, Mikkelsen TS, Rinn JL. The dynamics and regulators of cell fate decisions are revealed by pseudotemporal ordering of single cells. Nat Biotechnol. 2014;32(4):381–6. Epub 20140323. doi: 10.1038/nbt.2859. PubMed PMID: 24658644; PMCID: PMC4122333.

40. Shull LC, Lencer ES, Kim HM, Goyama S, Kurokawa M, Costello JC, Jones K, Artinger KB. PRDM paralogs antagonistically balance Wnt/β-catenin activity during craniofacial chondrocyte differentiation. Development. 2022;149(4). Epub 20220224. doi: 10.1242/dev.200082. PubMed PMID: 35132438; PMCID: PMC8918787.

41. Ramírez F, Ryan DP, Grüning B, Bhardwaj V, Kilpert F, Richter AS, Heyne S, Dündar F, Manke T. deepTools2: a next generation web server for deep-sequencing data analysis. Nucleic Acids Res. 2016;44(W1):W160–5. Epub 20160413. doi: 10.1093/nar/gkw257. PubMed PMID: 27079975; PMCID: PMC4987876.

42. Lopez-Delisle L, Rabbani L, Wolf J, Bhardwaj V, Backofen R, Grüning B, Ramírez F, Manke T. pyGenomeTracks: reproducible plots for multivariate genomic datasets. Bioinformatics. 2021;37(3):422–3. doi: 10.1093/bioinformatics/btaa692. PubMed PMID: 32745185; PMCID: PMC8058774.

43. Love MI, Huber W, Anders S. Moderated estimation of fold change and dispersion for RNA-seq data with DESeq2. Genome biology. 2014;15(12):550.

44. Fisch KM, Gamini R, Alvarez-Garcia O, Akagi R, Saito M, Muramatsu Y, Sasho T, Koziol JA, Su AI, Lotz MK. Identification of transcription factors responsible for dysregulated networks in human osteoarthritis cartilage by global gene expression analysis. Osteoarthritis Cartilage. 2018;26(11):1531–8. Epub 20180803. doi: 10.1016/j.joca.2018.07.012. PubMed PMID: 30081074; PMCID: PMC6245598.

45. Horn KH, Warner DR, Pisano M, Greene RM. PRDM16 expression in the developing mouse embryo. Acta Histochemica. 2011;113(2):150–5. doi: 10.1016/j.acthis.2009.09.006.

46. Long F, Linsenmayer TF. Regulation of growth region cartilage proliferation and differentiation by perichondrium. Development. 1998;125(6):1067–73. doi: 10.1242/dev.125.6.1067. PubMed PMID: 9463353.

47. Svandova E, Anthwal N, Tucker AS, Matalova E. Diverse Fate of an Enigmatic Structure: 200 Years of Meckel’s Cartilage. Front Cell Dev Biol. 2020;8:821. Epub 20200828. doi: 10.3389/fcell.2020.00821. PubMed PMID: 32984323; PMCID: PMC7484903.

48. Mizoguchi T, Ono N. The diverse origin of bone-forming osteoblasts. J Bone Miner Res. 2021;36(8):1432–47. Epub 20210712. doi: 10.1002/jbmr.4410. PubMed PMID: 34213032; PMCID: PMC8338797.

49. Ma HL, Blanchet TJ, Peluso D, Hopkins B, Morris EA, Glasson SS. Osteoarthritis severity is sex dependent in a surgical mouse model. Osteoarthritis Cartilage. 2007;15(6):695–700. Epub 20070103. doi: 10.1016/j.joca.2006.11.005. PubMed PMID: 17207643.

50. Colbath A, Haubruck P. Closing the gap: sex-related differences in osteoarthritis and the ongoing need for translational studies. Ann Transl Med. 2023;11(10):339. Epub 20230627. doi: 10.21037/atm-23-1546. PubMed PMID: 37675305; PMCID: PMC10477647.

51. Chuikov S, Levi BP, Smith ML, Morrison SJ. Prdm16 promotes stem cell maintenance in multiple tissues, partly by regulating oxidative stress. Nat Cell Biol. 2010;12(10):999–1006. Epub 20100912. doi: 10.1038/ncb2101. PubMed PMID: 20835244; PMCID: PMC2948585.

52. McGlynn KA, Sun R, Vonica A, Rudzinskas S, Zhang Y, Perkins AS. Prdm3 and Prdm16 cooperatively maintain hematopoiesis and clonogenic potential. Exp Hematol. 2020;85:20–32.e3. Epub 20200511. doi: 10.1016/j.exphem.2020.04.010. PubMed PMID: 32437910.

53. Gudmundsson KO, Nguyen N, Oakley K, Han Y, Gudmundsdottir B, Liu P, Tessarollo L, Jenkins NA, Copeland NG, Du Y. Prdm16 is a critical regulator of adult long-term hematopoietic stem cell quiescence. Proc Natl Acad Sci U S A. 2020;117(50):31945–53. Epub 20201202. doi: 10.1073/pnas.2017626117. PubMed PMID: 33268499; PMCID: PMC7749346.

54. Shimada IS, Acar M, Burgess RJ, Zhao Z, Morrison SJ. Prdm16 is required for the maintenance of neural stem cells in the postnatal forebrain and their differentiation into ependymal cells. Genes Dev. 2017;31(11):1134–46. Epub 20170711. doi: 10.1101/gad.291773.116. PubMed PMID: 28698301; PMCID: PMC5538436.

55. Seale P, Kajimura S, Yang W, Chin S, Rohas LM, Uldry M, Tavernier G, Langin D, Spiegelman BM. Transcriptional control of brown fat determination by PRDM16. Cell Metab. 2007;6(1):38–54. doi: 10.1016/j.cmet.2007.06.001. PubMed PMID: 17618855; PMCID: PMC2564846.

56. Jin Y, Cong Q, Gvozdenovic-Jeremic J, Hu J, Zhang Y, Terkeltaub R, Yang Y. Enpp1 inhibits ectopic joint calcification and maintains articular chondrocytes by repressing hedgehog signaling. Development. 2018;145(18). Epub 20180920. doi: 10.1242/dev.164830. PubMed PMID: 30111653; PMCID: PMC6176935.

57. Takahata Y, Hagino H, Kimura A, Urushizaki M, Kobayashi S, Wakamori K, Fujiwara C, Nakamura E, Yu K, Kiyonari H, Bando K, Murakami T, Komori T, Hata K, Nishimura R. Smoc1 and Smoc2 regulate bone formation as downstream molecules of Runx2. Commun Biol. 2021;4(1):1199. Epub 20211019. doi: 10.1038/s42003-021-02717-7. PubMed PMID: 34667264; PMCID: PMC8526618.

58. Abe M, Michikami I, Fukushi T, Abe A, Maeda Y, Ooshima T, Wakisaka S. Hand2 regulates chondrogenesis in vitro and in vivo. Bone. 2010;46(5):1359–68. Epub 20091122. doi: 10.1016/j.bone.2009.11.022. PubMed PMID: 19932774.

59. Arnold MA, Kim Y, Czubryt MP, Phan D, McAnally J, Qi X, Shelton JM, Richardson JA, Bassel-Duby R, Olson EN. MEF2C transcription factor controls chondrocyte hypertrophy and bone development. Dev Cell. 2007;12(3):377–89. doi: 10.1016/j.devcel.2007.02.004. PubMed PMID: 17336904.

60. Ruan X, Jin X, Sun F, Pi J, Jinghu Y, Lin X, Zhang N, Chen G. IGF signaling pathway in bone and cartilage development, homeostasis, and disease. Faseb j. 2024;38(17):e70031. doi: 10.1096/fj.202401298R. PubMed PMID: 39206513.

61. Clemmons DR, Busby WH, Jr., Garmong A, Schultz DR, Howell DS, Altman RD, Karr R. Inhibition of insulin-like growth factor binding protein 5 proteolysis in articular cartilage and joint fluid results in enhanced concentrations of insulin-like growth factor 1 and is associated with improved osteoarthritis. Arthritis Rheum. 2002;46(3):694–703. doi: 10.1002/art.10222. PubMed PMID: 11920405.

62. Gadomski SJ, Mui BWH, Gorodetsky R, Paravastu SS, Featherall J, Li L, Haffey A, Kim JC, Kuznetsov SA, Futrega K, Lazmi-Hailu A, Merling RK, Martin D, McCaskie AW, Robey PG. Time- and cell-specific activation of BMP signaling restrains chondrocyte hypertrophy. iScience. 2024;27(8):110537. Epub 20240718. doi: 10.1016/j.isci.2024.110537. PubMed PMID: 39193188; PMCID: PMC11347861.

63. Yano F, Ohba S, Murahashi Y, Tanaka S, Saito T, Chung UI. Runx1 contributes to articular cartilage maintenance by enhancement of cartilage matrix production and suppression of hypertrophic differentiation. Sci Rep. 2019;9(1):7666. Epub 20190521. doi: 10.1038/s41598-019-43948-3. PubMed PMID: 31113964; PMCID: PMC6529519.

