## Supplemental Figures for "PRDM16 Coordinates Genetic and Epigenetic Programs Governing Chondrogenesis and Chondrocyte Phenotype Specification in the Knee Joint"

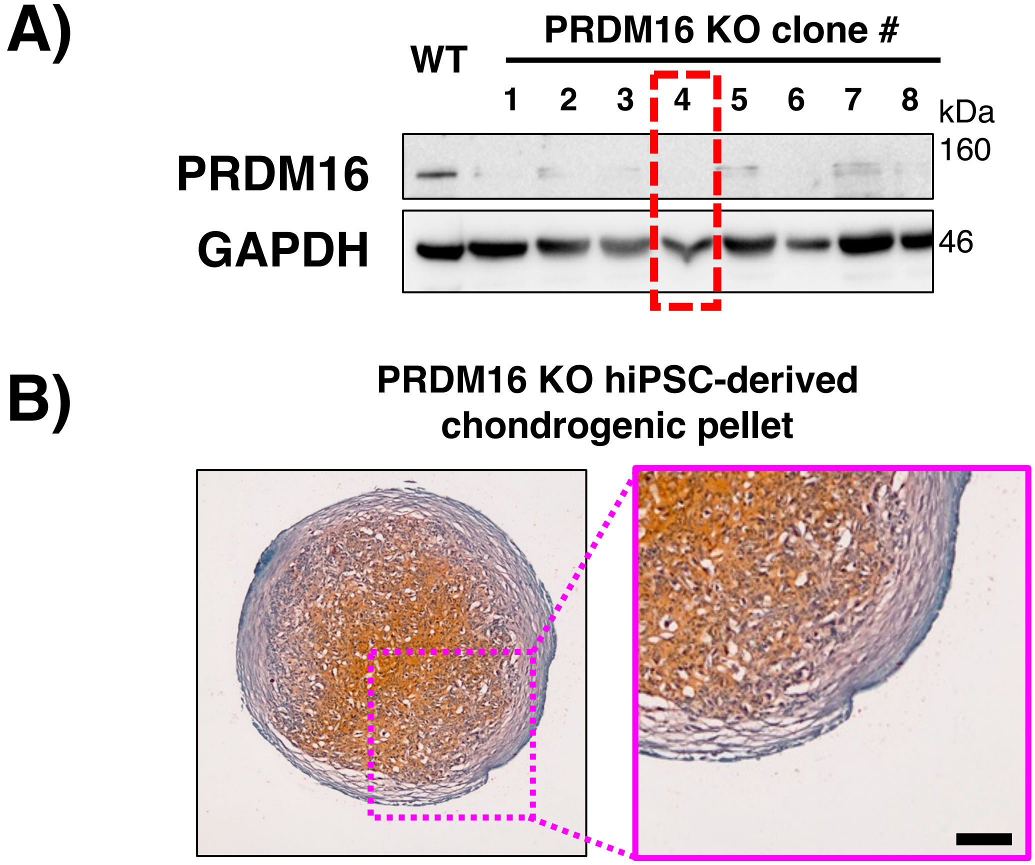


**Supp. Fig. 1: Clonal selection of PRDM16 KO.** (**A**) Representative Western Blot images of single-cell clonal selection of CRISPR/Cas9-mediated PRDM16 KO in hiPSCs. Note clone 4 (red box) was used for chondrogenesis and downstream CUT&RUN-seq. (**B**) Significantly inferior chondrogenic differentiation of PRDM16 KO hiPSCs when compared WT hiPSCs (see **Fig. 5B**). Black scale bar is 50$\mu$m.

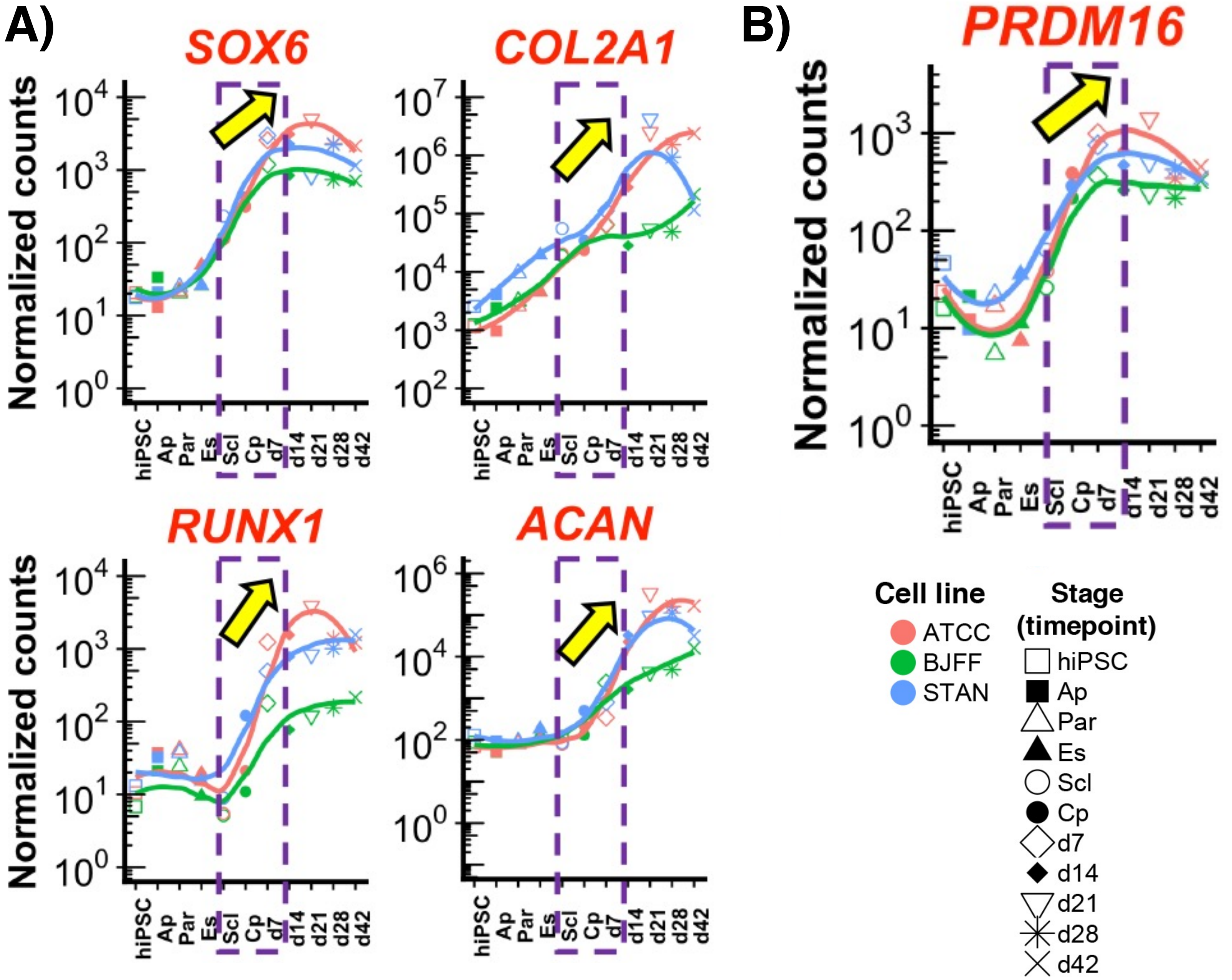


**Supp. Fig. 2: Bulk RNA-seq reveals that PRDM16 expression is increased across the $C_P$ stage and maintained for multiple days of chondrogenic cell culture.** (**A**) Upregulation of essential chondrogenic markers including *SOX6*, *COL2A1*, *RUNX1*, and *ACAN* during hiPSC chondrogenesis. (**B**) PRDM16 expression exhibited a positive association with these markers during hiPSC chondrogenesis. This expression profile was confirmed across 3 independent hiPSC lines (ATCC, BJFF, STAN). Ap = anterior primitive streak, Par = paraxial mesoderm, Es = early somite, Scl = sclerotome, Cp = chondroprogenitor stages.

| | **Control** | **KD** | **OE** |
|---|---|---|---|
| nFeature_RNA_max | <9000 | <9000 | <10000 |
| nFeature_RNA_min | >3000 | >2000 | >3000 |
| percent_mito | <15 | <15 | <15 |
| FindCluster resolution | 0.8 | 0.9 | 0.9 |

**Supp. Table 1:** Filtering parameters for Seurat objects of each sample prior to integration.

| | Cluster | Cell Number | Percent |
|---|---|---|---|
| Control | CONTROL_0 | 1052 | 27.706% |
| | CONTROL_1 | 683 | 17.988% |
| | CONTROL_2 | 461 | 12.141% |
| | CONTROL_3 | 618 | 16.276% |
| | CONTROL_4 | 122 | 3.213% |
| | CONTROL_5 | 204 | 5.373% |
| | CONTROL_6 | 280 | 7.374% |
| | CONTROL_7 | 154 | 4.056% |
| | CONTROL_8 | 142 | 3.740% |
| | CONTROL_9 | 37 | 0.974% |
| | CONTROL_10 | 44 | 1.159% |
| KD | KNOCKDOWN_0 | 751 | 21.587% |
| | KNOCKDOWN_1 | 838 | 24.087% |
| | KNOCKDOWN_2 | 343 | 9.859% |
| | KNOCKDOWN_3 | 562 | 16.154% |
| | KNOCKDOWN_4 | 165 | 4.743% |
| | KNOCKDOWN_5 | 355 | 10.204% |
| | KNOCKDOWN_6 | 101 | 2.903% |
| | KNOCKDOWN_7 | 126 | 3.622% |
| | KNOCKDOWN_8 | 90 | 2.587% |
| | KNOCKDOWN_9 | 118 | 3.392% |
| | KNOCKDOWN_10 | 30 | 0.862% |
| OE | OVEREXPRESSED_0 | 290 | 11.628% |
| | OVEREXPRESSED_1 | 175 | 7.017% |
| | OVEREXPRESSED_2 | 810 | 32.478% |
| | OVEREXPRESSED_3 | 201 | 8.059% |
| | OVEREXPRESSED_4 | 466 | 18.685% |
| | OVEREXPRESSED_5 | 92 | 3.689% |
| | OVEREXPRESSED_6 | 167 | 6.696% |
| | OVEREXPRESSED_7 | 97 | 3.889% |
| | OVEREXPRESSED_8 | 93 | 3.729% |
| | OVEREXPRESSED_9 | 13 | 0.521% |
| | OVEREXPRESSED_10 | 90 | 3.609% |

**Supp. Table. 2:** Cell number and percent cells in each cluster per treatment group. This table is associated with the UMAP in Fig. 5D.

| Group | Cluster | Cell Number | Percent |
|---|---|---|---|
| Control | CONTROL_0 | 910 | 31.401% |
| | CONTROL_1 | 968 | 33.402% |
| | CONTROL_2 | 542 | 18.703% |
| | CONTROL_3 | 203 | 7.005% |
| | CONTROL_4 | 275 | 9.489% |
| KD | KNOCKDOWN_0 | 1061 | 39.516% |
| | KNOCKDOWN_1 | 705 | 26.257% |
| | KNOCKDOWN_2 | 446 | 16.611% |
| | KNOCKDOWN_3 | 361 | 13.445% |
| | KNOCKDOWN_4 | 112 | 4.171% |
| OE | OVEREXPRESSED_0 | 565 | 29.442% |
| | OVEREXPRESSED_1 | 905 | 47.160% |
| | OVEREXPRESSED_2 | 183 | 9.536% |
| | OVEREXPRESSED_3 | 92 | 4.794% |
| | OVEREXPRESSED_4 | 174 | 9.067% |

**Supp. Table. 3:** Frequency and percent cells in subset data of each cluster per treatment group. This table is associated with the UMAP in Fig. 5E.

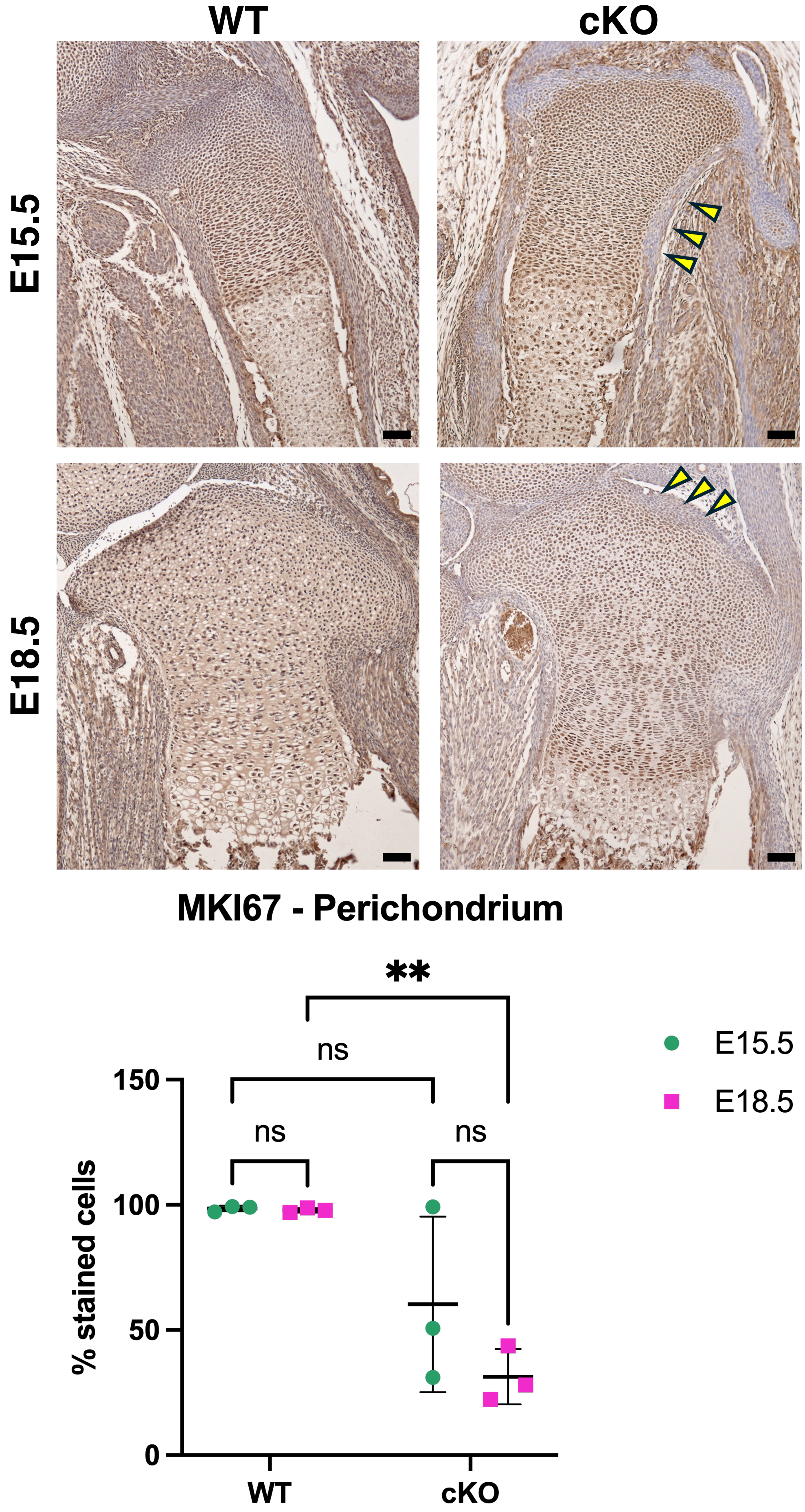


**Supp. Fig. 3: Embryonic cKO mice exhibit reduced MKI67 expression in the perichondrium at E18.5.** (**A**) Histological staining of MKI67 and (**B**) quantification thereof, scale bar = 50μm. Yellow arrow-heads: perichondrium. One-Way ANOVA, Tukey's Post-hoc (**p < 0.01).

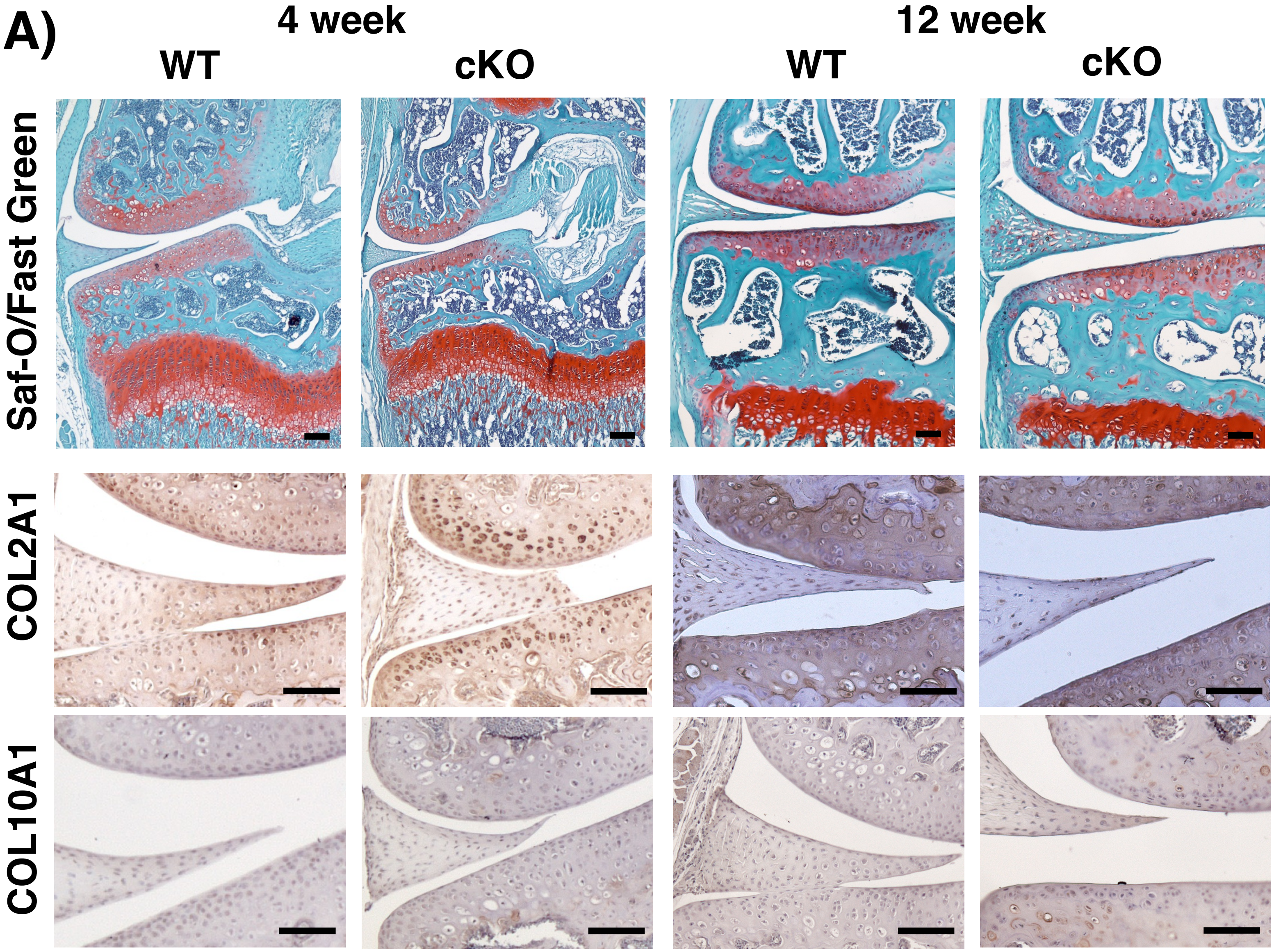


**Supp. Fig. 4: Physiological evaluation of 4- or 12-wk-old PRDM16 WT and cKO female mice cartilage.** (**A**) Histological staining of knee joints: Safranin-O Fast Green, Type II Collagen, and Type 10 Collagen. Black scale bar = 50$\mu$m. Pink arrowhead indicates unstained cells. (**B**) Weights of mice at day of sacrifice. ns = nonsignificant.

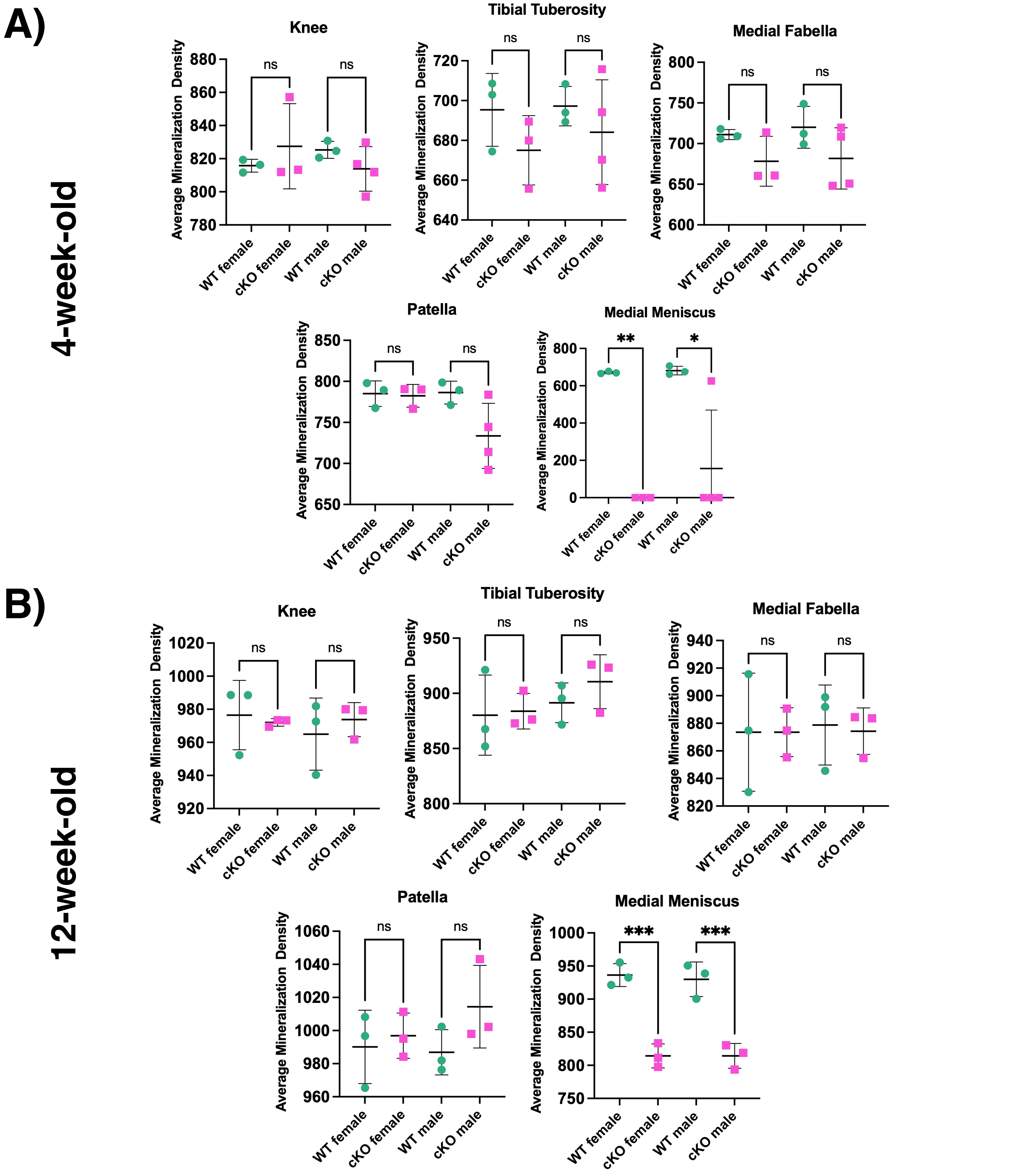


**Supp. Fig. 5:** *μ*CT analysis of (**A**) 4-wk-old and **(B**) 12-wk-old mouse bone mineralization density reveals reduced bone mineralization density (mg HA/cm³) of medial meniscus in cKO mice vs. WT. Data analyzed by Two-Way ANOVA, Tukey's post-hoc (*p < 0.05. **p < 0.01, ***p < 0.001).

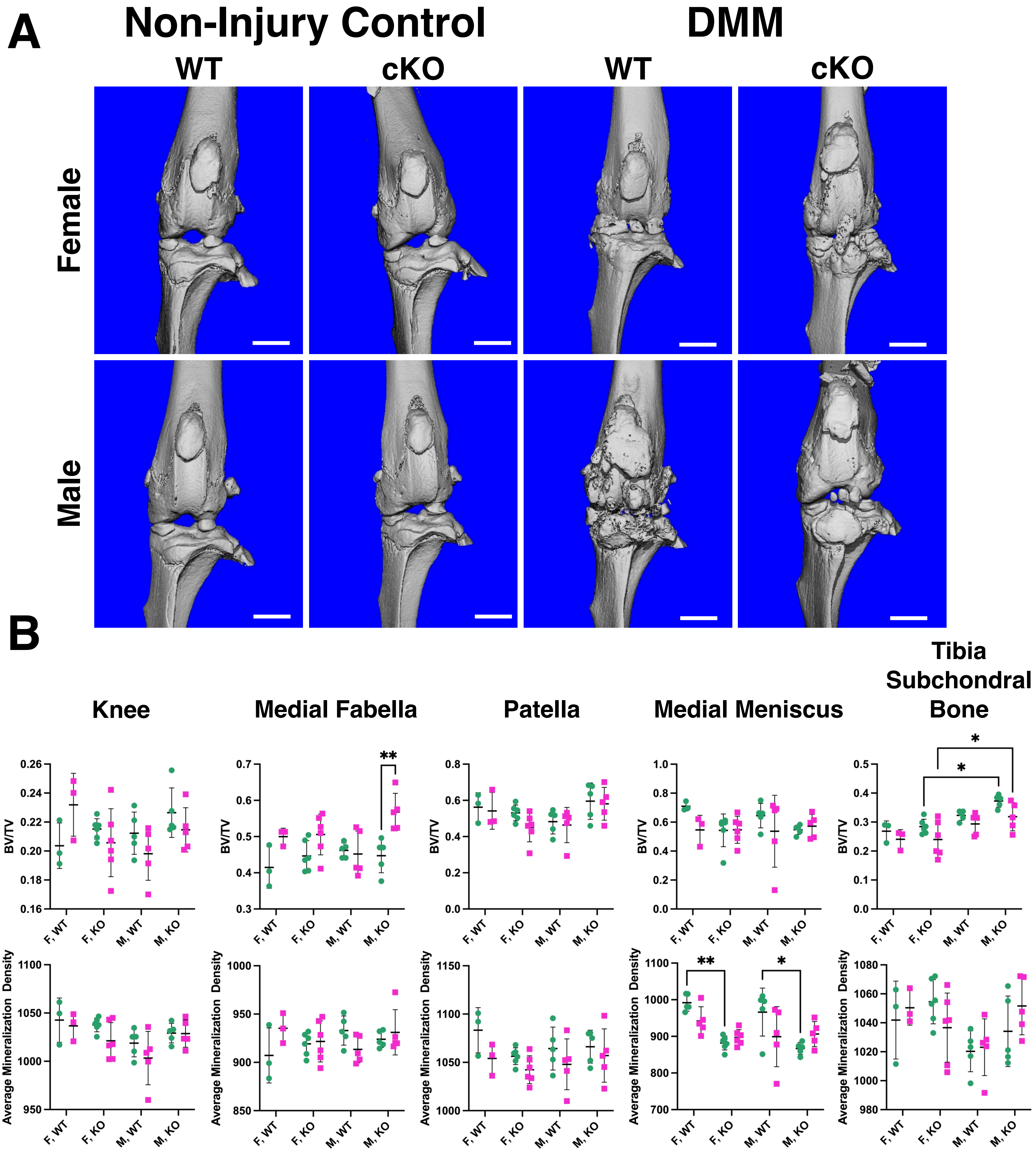


**Supp. Fig. 6**: **DMM injury model reveals reduced bone quality and poor joint homeostasis in cKO mice.** (**A**) $\mu$CT images of male WT and cKO non-surgery (left) and DMM (right) joints. (**B**) BV/TV and Average Mineralization Density (mg HA/cm$^3$) in select ROIs. Data analyzed by Two Way ANOVA, Tukey's post-hoc, $^*p < 0.05$.

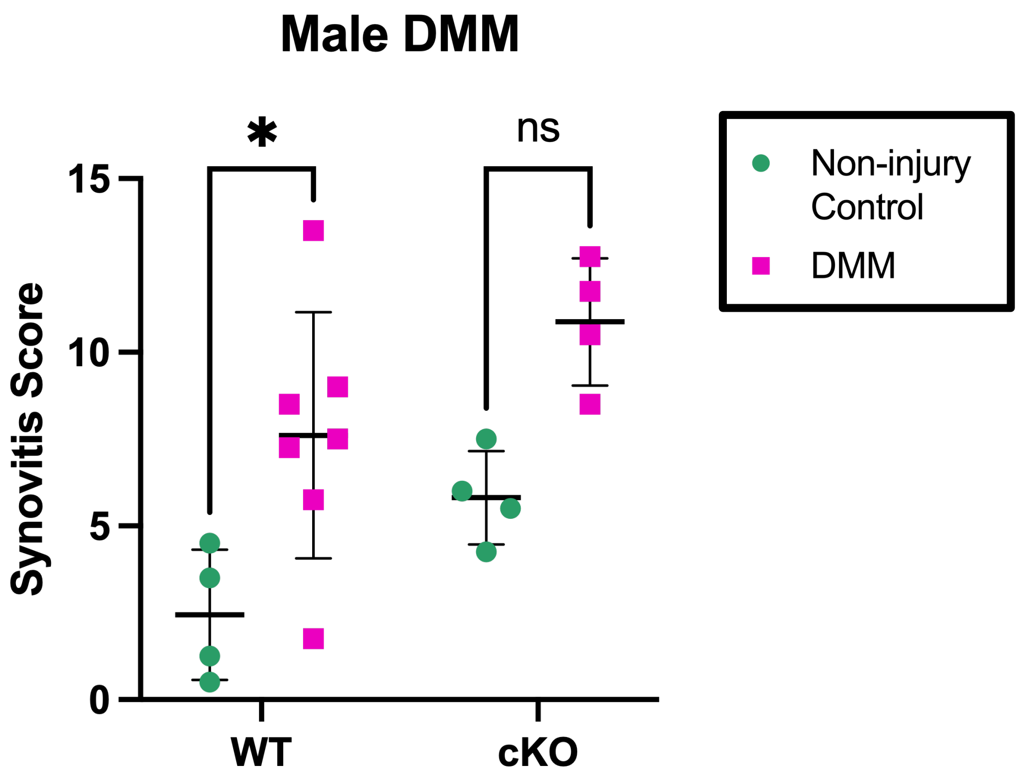


**Supp. Fig. 7: Synovitis score of male DMM and non-injury control joints.** Only male mice were analyzed and quantified because DMM joints of female mice exhibited minimal cartilage degradation. Data analyzed by Two-Way ANOVA, Tukey’s post-hoc, *$p < 0.05$.

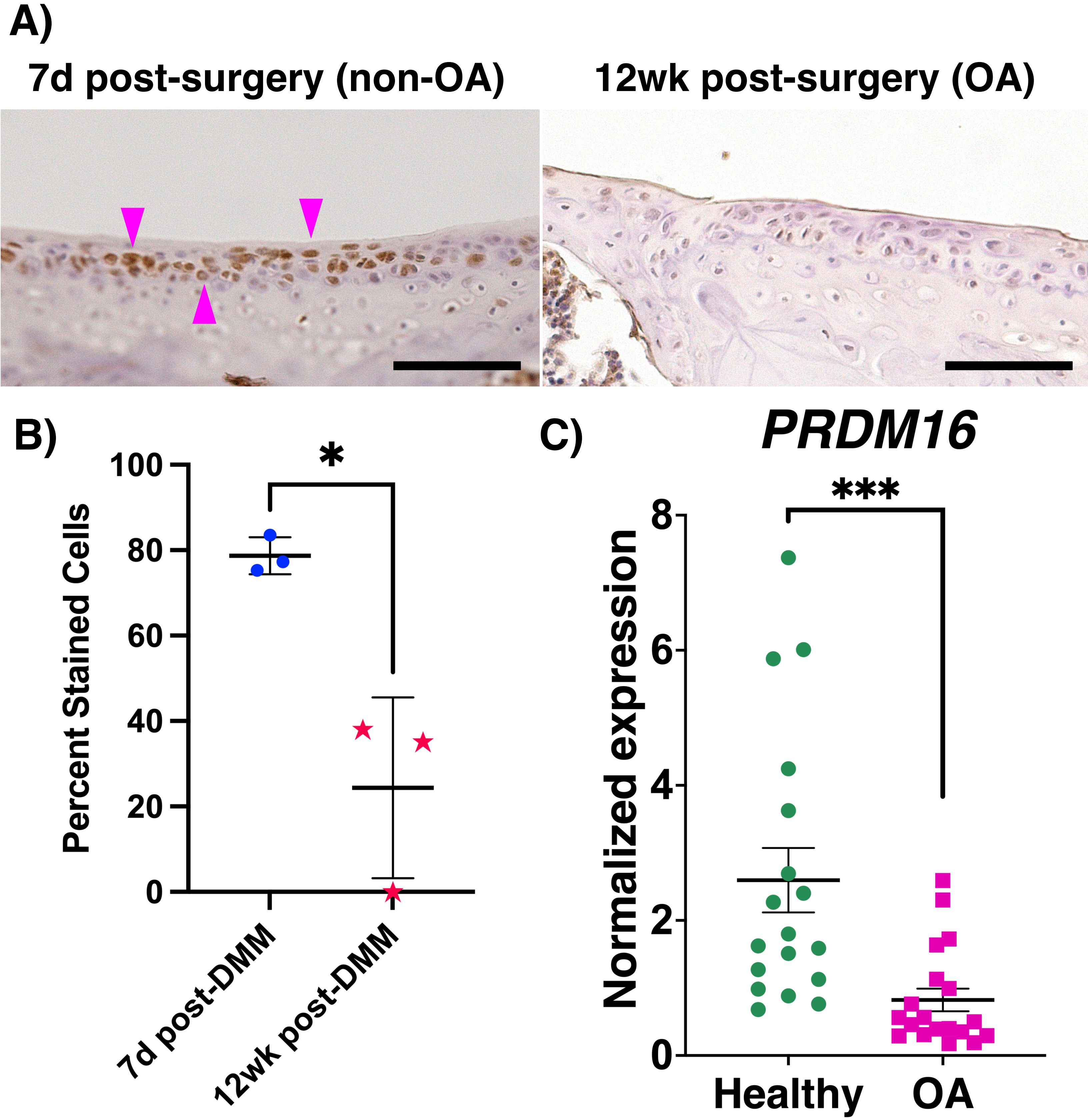
A)
7d post-surgery (non-OA)
12wk post-surgery (OA)
B)
Percent Stained Cells
100
80
60
40
20
0
*
7d post-DMM
12wk post-DMM
C)
PRDM16
Normalized expression
8
6
4
2
0
***
Healthy
OA

**Supp. Fig. 8: C57BL/6J mice with DMM exhibit reduced PRDM16 expression in the uncalcified articular cartilage at 12-weeks post-injury.** (**A**) Histological staining of PRDM16 in tibia of injured joints. Black scale bar = 50*μ*m. Pink arrowhead indicates $PRDM16^+$ cells. (**B**) Quantification of percent $PRDM16^+$ cells in the uncalcified tibial articular cartilage. (**C**) Human OA cartilage has significantly reduced PRDM16 expression compared to Healthy. Publicly available dataset (Fisch+, 2018). Student's t-test (*$p < 0.05$, ***$p < 0.001$).

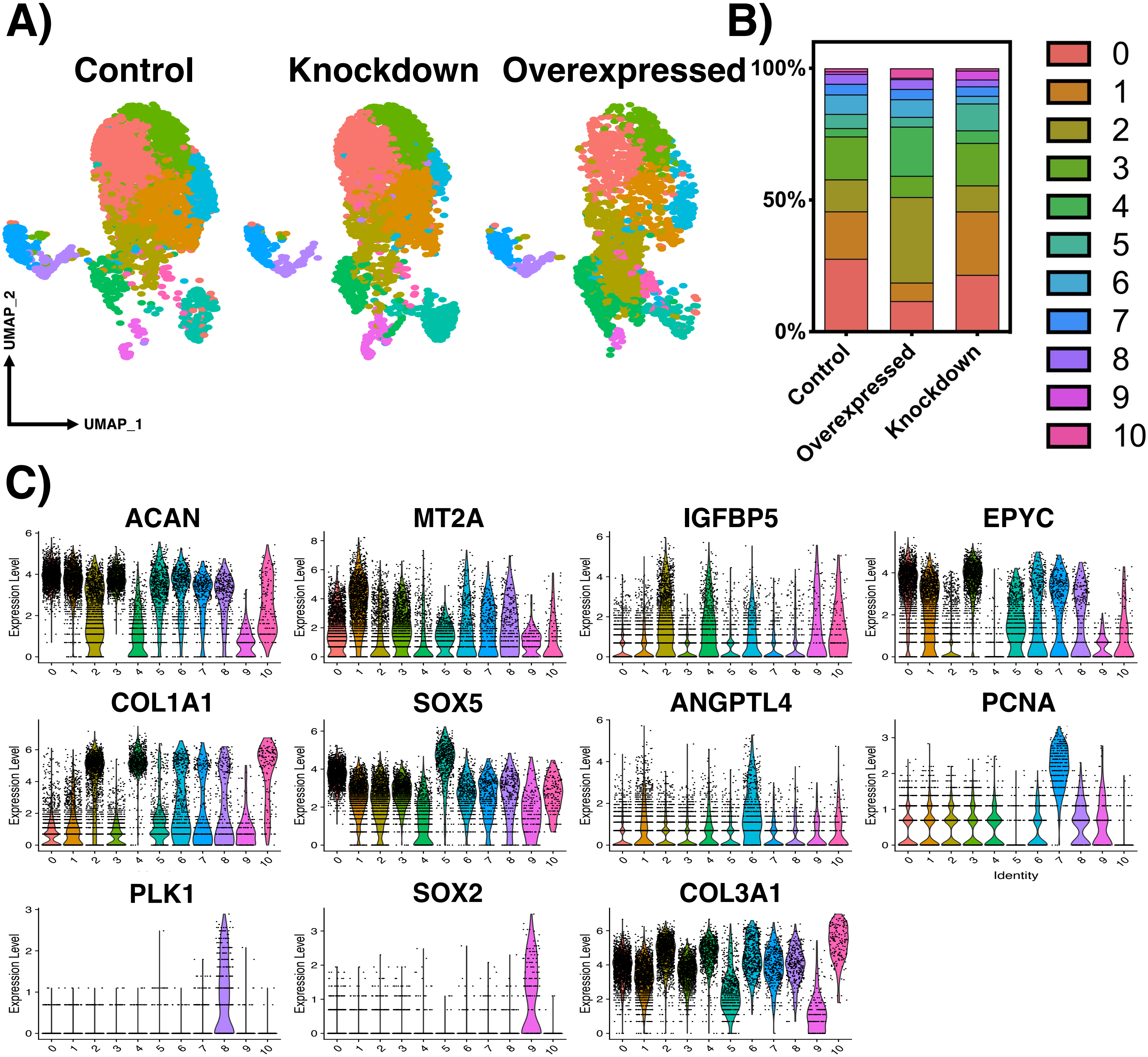


**Supp. Fig. 9:** (**A**) 11 unique populations are conserved across all treatment groups. (**B**) Percentage of cells located in each chondrocyte cluster per treatment group. (**C**) Select marker genes for each cluster and correlating expression among all other clusters.

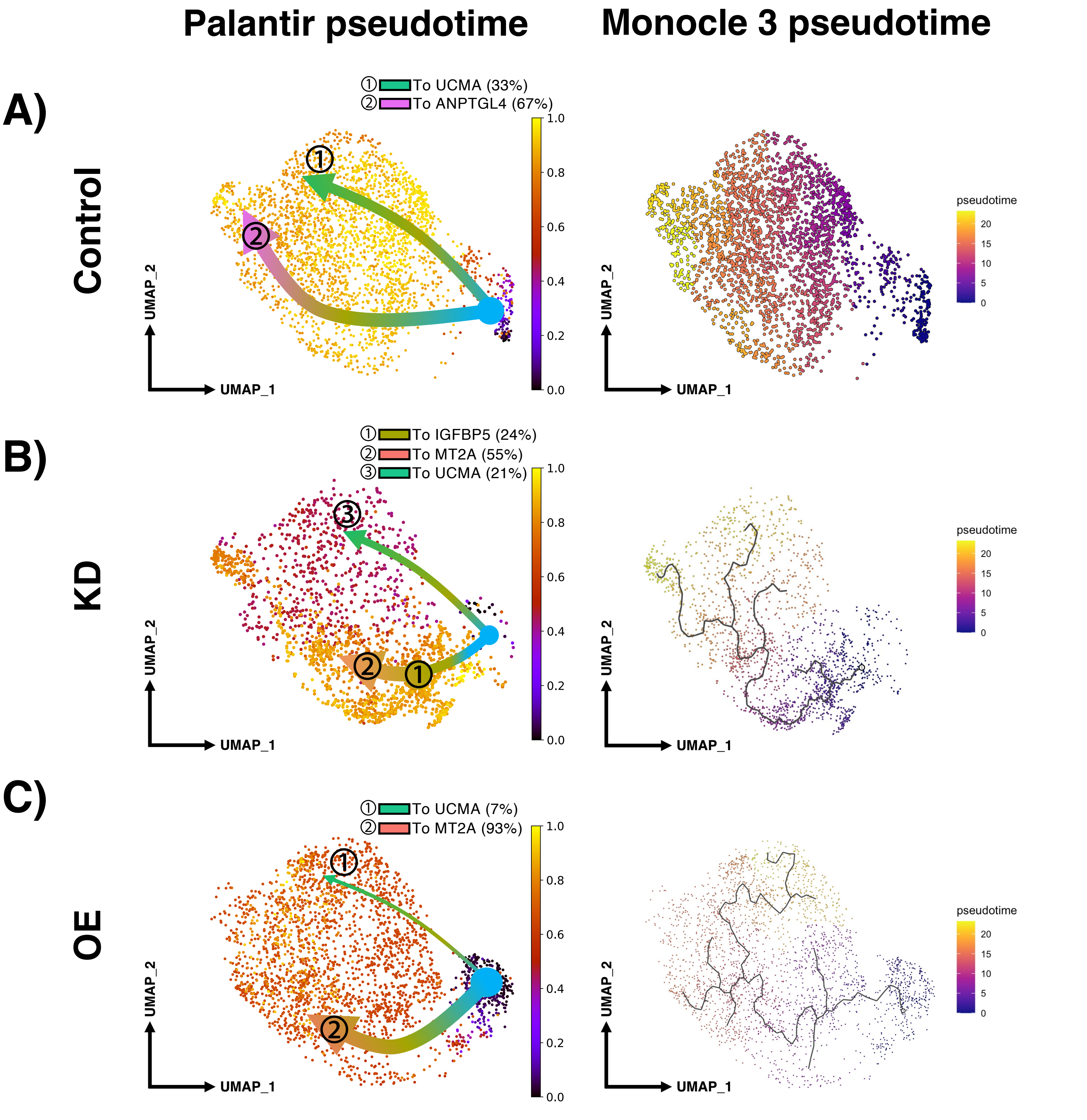


**Supp. Fig. 10: Palantir and Monocle 3 pseudotime analyses** for (**A**) Control, (**B**) KD, and (**C**) OE chondrogenic pellets. Similar to RNA velocity and random walk (**Fig. 6**), both Palantir and Monocle 3 pseudotime algorithms predict the SOX5$^{high}$/SOX6$^{high}$ chondroprogenitor population as the initial state of the differentiation trajectory.

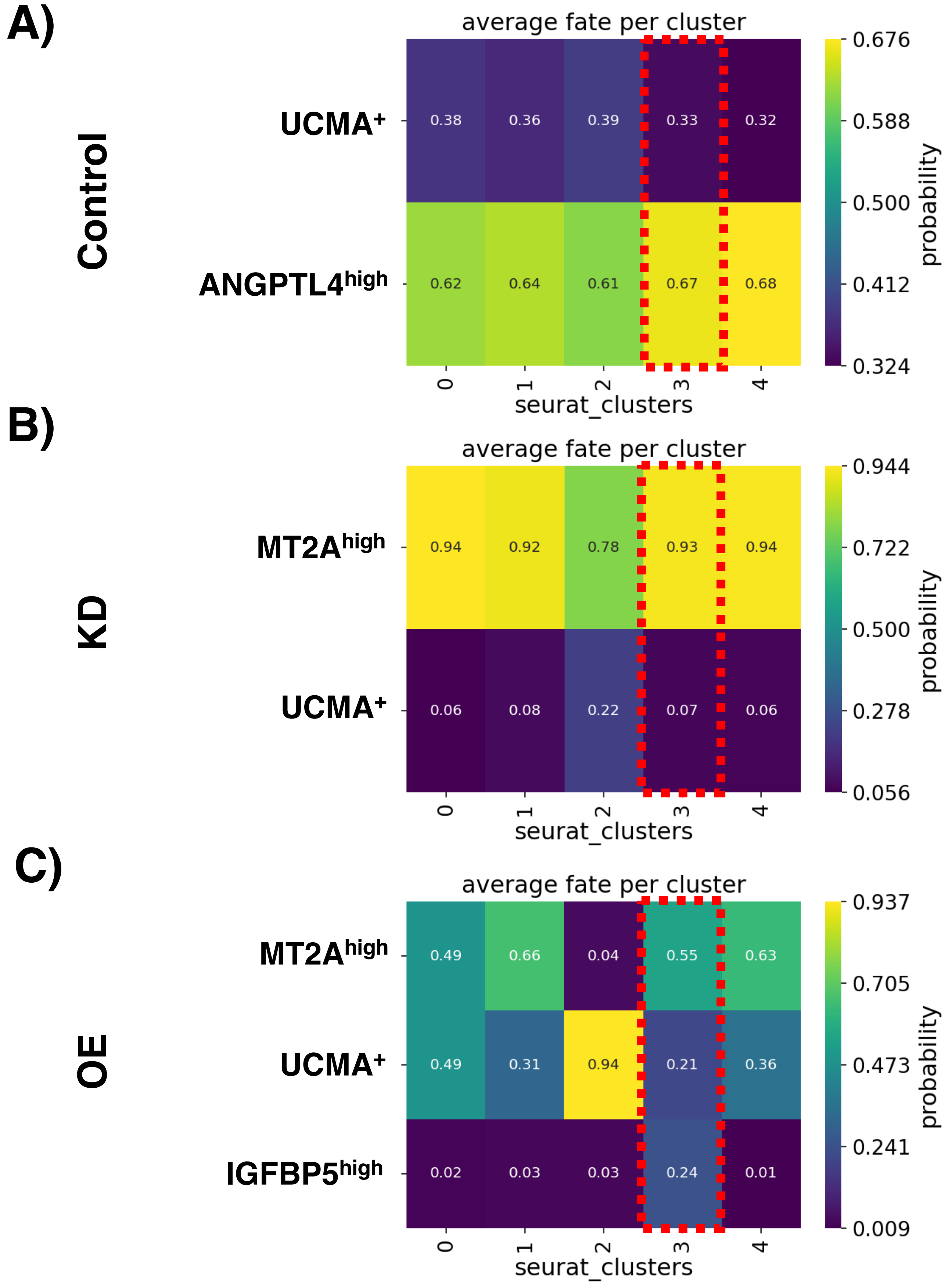


**Supp. Fig. 11: Average fate probability per cell cluster computed by CellRank 2** for (**A**) Control, (**B**) Knockdown, and (**C**) Overexpressed chondrogenic pellets. Seurat clusters 0: MT2A$^{high}$ chondrocytes, 1: IGFPB5$^{high}$ chondrocytes, 2: UCMA+ chondrocytes, 3: SOX5$^{high}$/SOX6$^{high}$ chondroprogenitors (red box), and 4: ANGPTL4$^{high}$ chondrocytes.

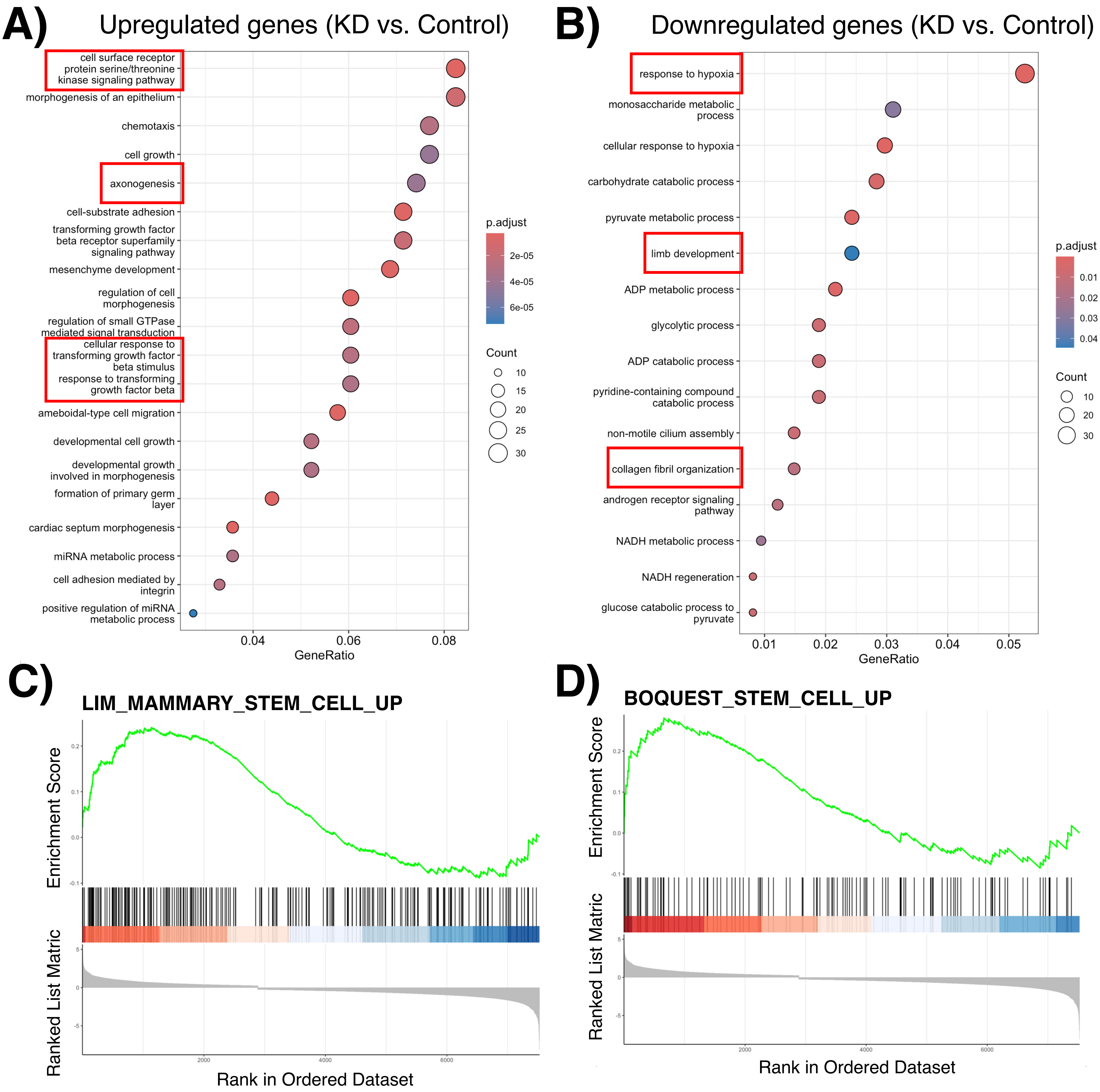
A)
Upregulated genes (KD vs. Control)
cell surface receptor protein serine/threonine kinase signaling pathway
morphogenesis of an epithelium
chemotaxis
cell growth
axonogenesis
cell-substrate adhesion
transforming growth factor beta receptor superfamily signaling pathway
mesenchyme development
regulation of cell morphogenesis
regulation of small GTPase mediated signal transduction
cellular response to transforming growth factor beta stimulus
response to transforming growth factor beta
ameboidal-type cell migration
developmental cell growth
developmental growth involved in morphogenesis
formation of primary germ layer
cardiac septum morphogenesis
miRNA metabolic process
cell adhesion mediated by integrin
positive regulation of miRNA metabolic process
0.04
0.06
0.08
GeneRatio
p.adjust
2e-05
4e-05
6e-05
Count
10
15
20
25
30
B)
Downregulated genes (KD vs. Control)
response to hypoxia
monosaccharide metabolic process
cellular response to hypoxia
carbohydrate catabolic process
pyruvate metabolic process
limb development
ADP metabolic process
glycolytic process
ADP catabolic process
pyridine-containing compound catabolic process
non-motile cilium assembly
collagen fibril organization
androgen receptor signaling pathway
NADH metabolic process
NADH regeneration
glucose catabolic process to pyruvate
0.01
0.02
0.03
0.04
0.05
GeneRatio
p.adjust
0.01
0.02
0.03
0.04
Count
10
20
30
C)
LIM_MAMMARY_STEM_CELL_UP
Enrichment Score
Ranked List Matric
Rank in Ordered Dataset
D)
BOQUEST_STEM_CELL_UP
Enrichment Score
Ranked List Matric
Rank in Ordered Dataset

**Supp. Fig. 12: GO functional and GSEA analyses reveal altered biological processes but increased stemness in SOX5$^{high}$/SOX6$^{high}$ chondroprogenitors in cells with KD PRDM16 vs. Control.** DEGs of SOX5$^{high}$/SOX6$^{high}$ chondroprogenitors in KD vs. Control were used for GO functional analysis using *clusterProfiler* R package . SOX5$^{high}$/SOX6$^{high}$ chondroprogenitors in cells with KD PRDM16 exhibited (**A**) up-regulation of GO terms related to Serine/threonine kinase signaling pathway, axonogenesis, TGF-β signaling pathway, etc. and (**B**) down-regulation of GO terms related to response to hypoxia, limb development, collagen fibril organization, and carbohydrate and glycolytic processes. Gene set enrichment analysis (GSEA) shows that gene sets (**C**) LIM_MAMMARY_STEM_CELL_UP and (**D**) BOQUEST_STEM_CELL_UP were identified in the SOX5$^{high}$/SOX6$^{high}$ chondroprogenitor population among KD cells.

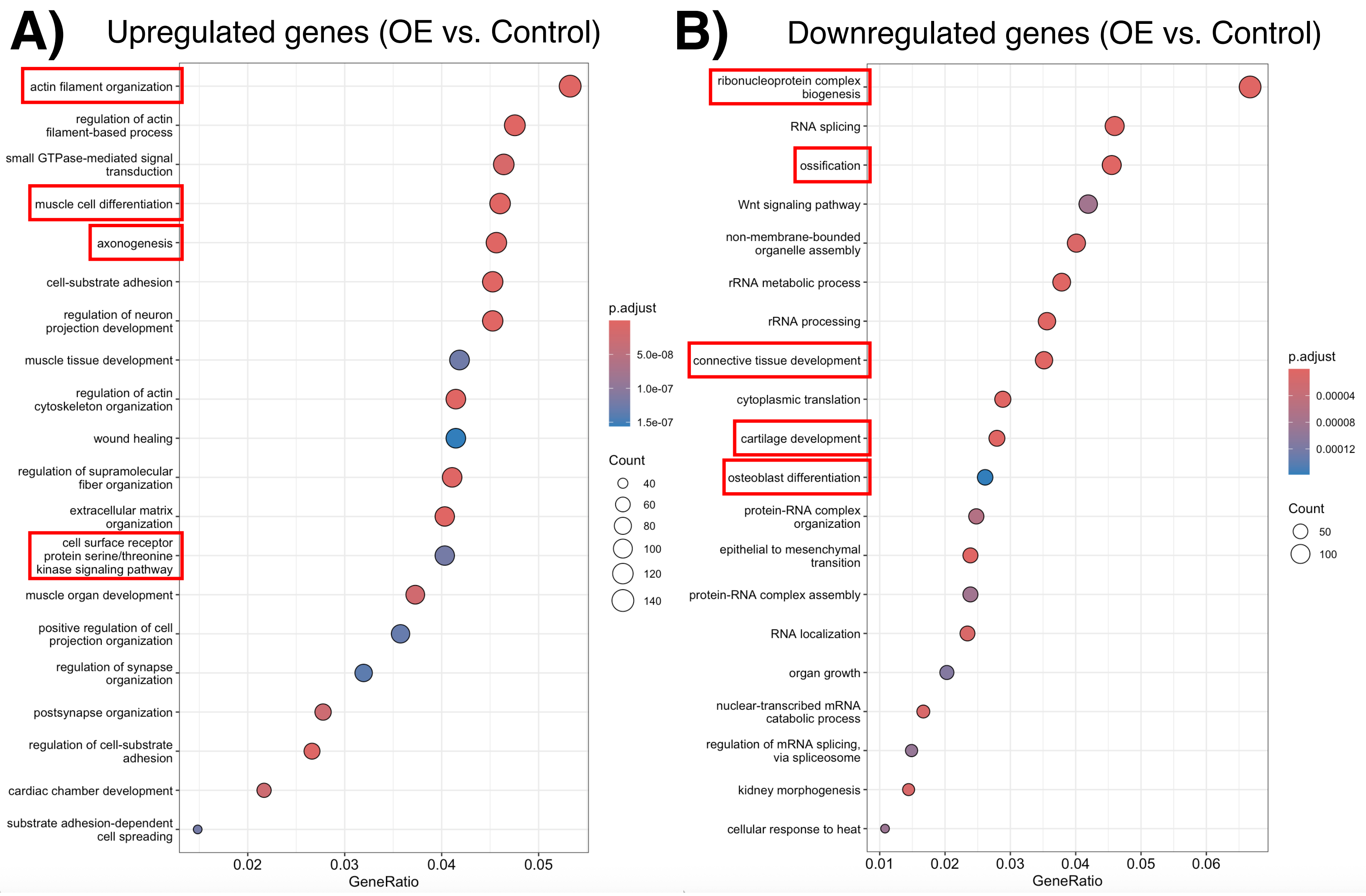


**Supp. Fig. 13: GO functional analysis reveals altered biological processes in IGFBP5$^{high}$ chondrocytes in cells with OE PRDM16 vs. Control.** DEGs of IGFBP5$^{high}$ chondrocytes in OE vs. Control were used for GO functional analysis using *clusterProfiler* R package . IGFBP5$^{high}$ chondrocytes in cells with OE PRDM16 exhibited (**A**) up-regulation of GO terms related to actin filament organization, axonogenesis, muscle cell differentiation, etc. and (**B**) down-regulation of GO terms related to ossification, cartilage development, osteoblast differentiation, connective tissue development, and several terms associated with rRNA biogenesis and processing.

| GO Term # | Biological Process |
| --- | --- |
| GO:0051216 | cartilage development |
| GO:0002062 | chondrocyte differentiation |
| GO:0060350 | cartilage cell differentiation |
| GO:0061035 | regulation of cartilage development |
| GO:0061036 | positive regulation of cartilage development |
| GO:0061037 | negative regulation of cartilage development |
| GO:0061975 | chondrocyte proliferation |
| GO:0030198 | extracellular matrix organization |
| GO:0032963 | collagen fibril organization |
| GO:0060452 | cartilage condensation |
| GO:0061973 | chondrocyte hypertrophy |
| GO:0003418 | growth plate cartilage development |

**Supp. Table 4:** Selected Cartilage-Related GO Terms.
